# Cellular and computational models reveal environmental and genetic interactions in *MMUT*-type methylmalonic aciduria

**DOI:** 10.1101/2022.08.10.503435

**Authors:** Charlotte Ramon, Florian Traversi, Céline Bürer, D. Sean Froese, Jörg Stelling

## Abstract

MMUT-type methylmalonic aciduria is a rare inherited metabolic disease caused by the loss of function of the methylmalonyl-CoA mutase (MMUT) enzyme. Patients develop symptoms resembling those of primary mitochondrial disorders, but the underlying causes of mitochondrial dysfunction remain unclear. Here, we examined environmental and genetic interactions in MMUT deficiency using a combination of computational modeling and cellular models to decipher pathways interacting with MMUT. Immortalized fibroblast (hTERT BJ5ta) MMUT-KO (MUTKO) clones displayed a mild mitochondrial impairment in standard glucose-based medium, but they did not to show increased reliance on respiratory metabolism nor reduced growth or viability. Consistently, our modeling predicted MUTKO specific growth phenotypes only for lower extracellular glutamine concentrations. Indeed, two of three MMUT-deficient BJ5ta cell lines showed a reduced viability in glutamine-free medium. Further, growth on 183 different carbon and nitrogen substrates identified increased NADH (nicotinamide adenine dinucleotide) metabolism of BJ5ta and HEK293 MUTKO cells compared to controls on purine- and glutamine-based substrates. With this knowledge, our modeling predicted 13 reactions interacting with MMUT that potentiate an effect on growth, primarily those of secondary oxidation of propionyl-CoA, oxidative phosphorylation and oxygen diffusion. Of these, we validated 3-hydroxyisobutytyl-CoA hydrolase (HIBCH) in the secondary propionyl-CoA oxidation pathway. Altogether, these results suggest compensation for the loss of MMUT function by increasing anaplerosis through glutamine or by diverting flux away from MMUT through the secondary propionyl-CoA oxidation pathway, which may have therapeutic relevance.

**sentence take-home message:** By perturbing metabolic pathways through genetic and environmental interventions in cellular and computational models of *MMUT*-type methylmalonic aciduria, we identified glutamine and secondary oxidative propionyl-CoA oxidation pathways as being important in the disease.

## Introduction

Isolated methylmalonic aciduria (MMAuria) is an inborn error of metabolism caused by the loss of function of the enzyme methylmalonyl coenzyme A mutase (MMUT), which transforms L-methylmalonyl-CoA into succinyl-CoA (metabolic reaction MMMm; Fig. 1). MMAuria may result from bi-allelic deleterious variants in the *MMUT* gene or in genes involved in the production of adenosyl-cobalamin, the cofactor of MMUT. Upon loss-of-function of the MMUT enzyme, upstream metabolites such as methylmalonic acid (MMA) and propionyl-CoA, as well as the latter’s derivatives propionyl-carnitine and 2-methylcitrate accumulate. Individuals affected by MMAuria may have life threatening episodes of metabolic crises and metabolic acidosis^1^. Despite therapies including a low protein diet and supplemental carnitine^2,3^ or even liver transplantation^4^, in the long-term, patients typically develop neurological complications and kidney failure^5,6^.

**Figure 1.**
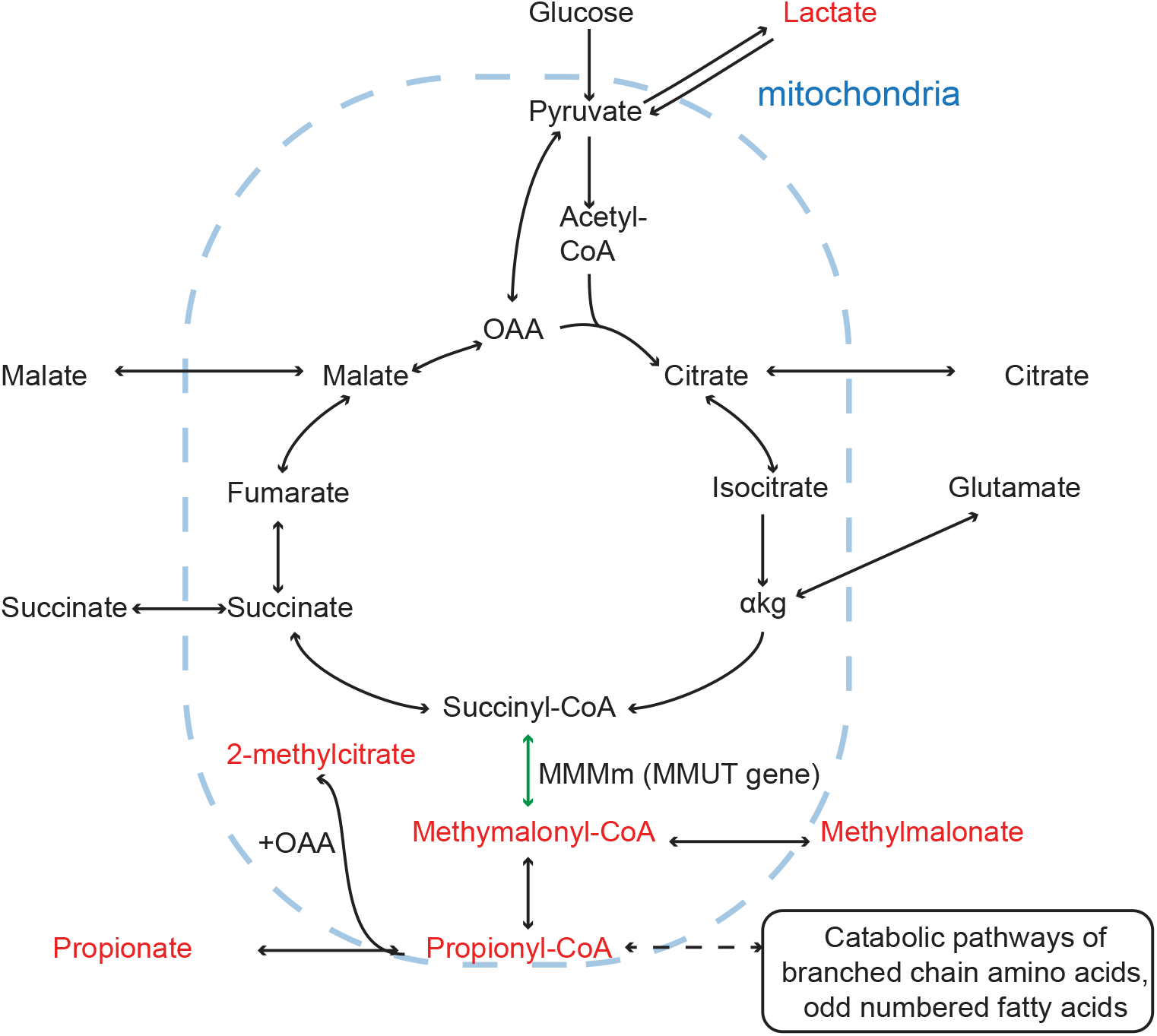
Metabolic pathways surrounding the MMUT reaction. MMUT catalyzes the reversible transformation of succinyl-CoA into methylmalonyl-CoA (green) in the mitochondria. Metabolites known to accumulate in patients affected by MMUT deficiency are colored in red. OAA: oxaloacetate. *α*kg: alpha-ketoglutarate.

Previous work has suggested neurological damage is associated with mitochondrial energy impairment^7^. This is supported by elevated levels of lactic acid in the globus pallidus, a region of the brain frequently affected, cerebrospinal fluid^8^ and plasma of patients^1,9^. Further, lower activities of cytochrome c oxidase in the liver, kidney and muscle of patients^10–12^ and in the liver of mutant mice^11^ have been reported. However, the activity of the other respiratory chain complexes did not show consistent results between different patients and studies^10–12^, for reasons that remain unclear^13^.

Two main, mutually non-exclusive hypotheses aim to explain the clinical and biochemical observations of MMAuria. The “toxic metabolite” hypothesis^14^ attributes the reduced activity of the tricarboxylic acid (TCA) cycle enzymes and respiratory complexes to inhibition by organic acids (e.g. MMA) and other compounds such as CoA metabolites (e.g. propionyl-CoA) that accumulate in the disease (Fig. 1). Indeed, MMA induces cell damage in neuronal cell culture systems^15^. MMA was shown to inhibit TCA cycle and oxidative phosphorylation associated enzymes (e.g. pyruvate carboxylase^16^) and transporters (e.g. malate shuttle^17^ and succinate transport ^18^) across mitochondrial membranes. However, MMA failed to directly inhibit respiratory complexes in submitochondrial particles from bovine heart, in particular complex II^19^. It was therefore suggested that other MMAuria associated metabolites might play a role and explain neuronal cell death, such as 2-methylcitrate or propionyl-CoA; they accumulate in neurons exposed to MMA^19^ and in patient plasma. Consistently, 2-methylcitrate was shown to inhibit enzymes of the TCA cycle^20^, and propionyl-CoA to inhibit pyruvate dehydrogenase^21^.The second, “TCA cycle depletion”, hypothesis suggests that the loss of MMUT enzyme function, which catalyzes an anaplerotic reaction replenishing the TCA cycle, leads to reduced TCA cycle flux and possible depletion of its metabolites^22^. This is consistent with succinate partially rescuing MMA-exposed neurons from death^19^.

To identify pathomechanisms in the network context, here we perturbed metabolic pathways through genetic and environmental interventions in cellular and computational models of MMUT-type MMAuria. We found increased NADH production of purine and glutamine metabolic pathways in MMUT-deficient cells, along with a faster cell death in glutamine-free medium. We also identified an interaction between HIBCH and MMUT enzyme. These results suggest that MMUT-deficient cells compensate for the loss of MMUT function through glutamine and secondary propionyl-CoA oxidation pathways.

## Results

### Establishment and characterization of a MMUT-deficient cell-line model

To avoid variation typical for cell lines obtained from different individuals such as patient fibroblasts, we created three MMUT deficient (MUTKO) cell lines from wild-type (WT) immortalized fibroblasts (BJ5tA) using CRISPR-Cas9. We isolated the mutant cell lines as single clones, each of which contained a homozygous frameshift mutation in exon 3 of the *MMUT* gene. Two separate clones harboured a deletion of two base pairs (c.613-614del, p(.Glu205Thrfs*5); MUTKO2 and MUTKO3) and one clone harbored an addition of one base pair (c.417dup, p.(Leu140Serfs*8); MUTKO7) (Supplementary figure 1). In all three mutated cell lines, we detected no full-length MMUT protein (Fig. 2 **a**), and no MMUT activity (Fig. 2 **b**). We checked for potential off-target effects of the CRISPR-Cas9 procedure by whole-genome sequencing. MUTKO-2, 3 and, 7 showed 6962, 5952 and 7098 novel variants compared to the parental cell line, respectively (Supplementary table 10). However, only 10-13 variants per clone potentially affected genes, mostly pseudogenes or genes of unknown function, and all variants were found on a single allele and were non-coding (intronic, upstream or downstream regions). Hence, the mutant cell lines have the desired metabolic perturbation and no other variants with a plausible effect on the metabolic phenotype.

**Figure 2.**
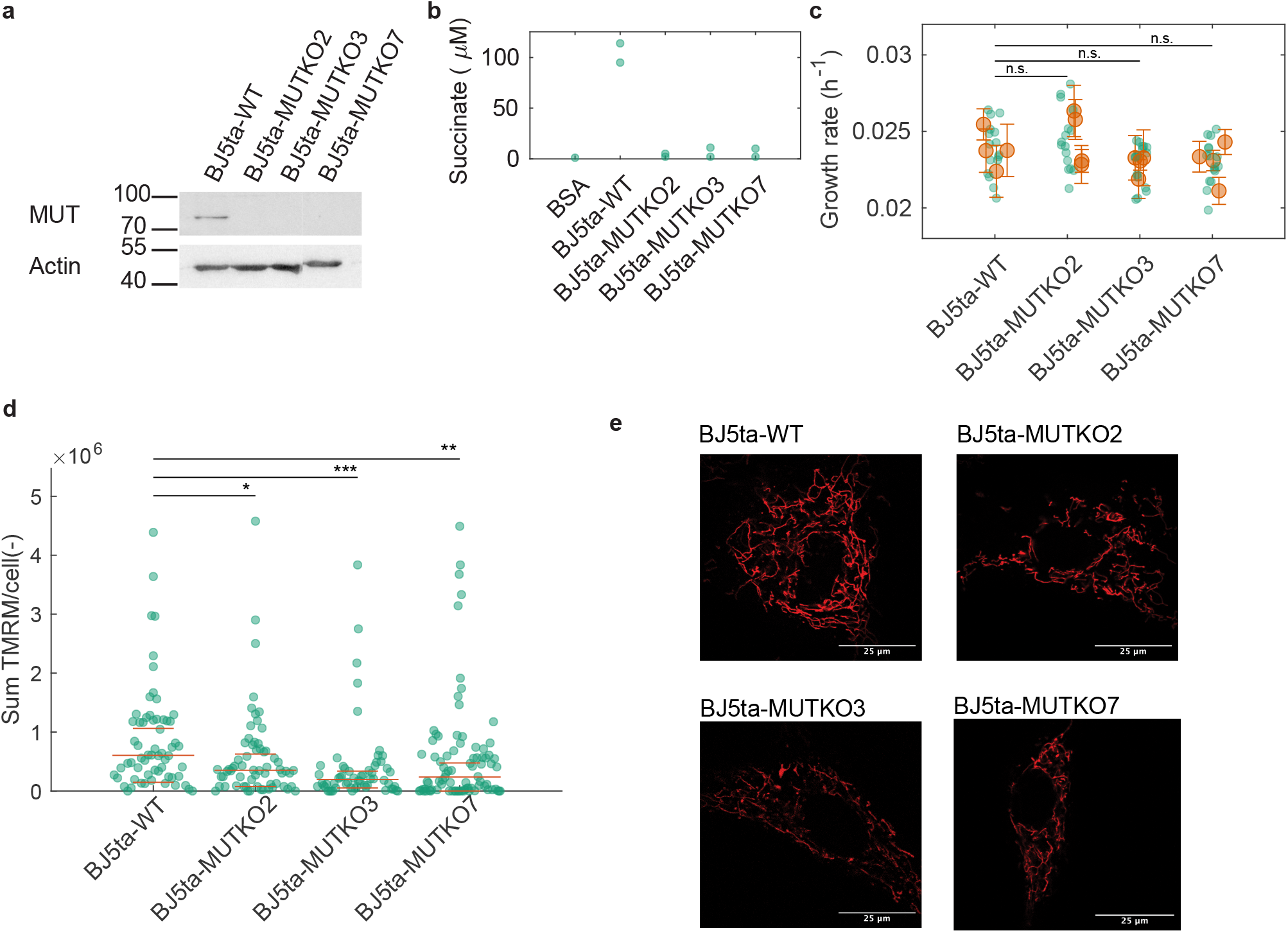
Characterization of MMUT-deficient cell lines. **a** Western blot validating the presence or absence of MMUT in the mutant cell lines. Actin served as loading control. **b** MMUT activity assay. Methylmalonyl-CoA was transformed into succinyl-CoA using cell lysates. Hydrolysis transforms succinyl-CoA into succinate, which is measured using mass spectrometry. BSA was used as negative control. **c** Growth rate of BJ5ta fibroblasts in the maintenance medium (DMEM mixed with M199, see **Methods**). Green dots: technical replicate; orange dots and bars: mean and standard deviation of the technical replicates for independent experiments. The overall means per mutated cell line were compared to WT, after adjusting for date effects. P-values were non-significant (n.s., *P − value >* 0.05) after correcting for multiple comparisons using Dunnett’s method. **d** Sum of TMRM fluorescence per cell. Points: sums of the blanked fluorescence of one cell (see **Methods**); wide red lines: median; narrow red lines: median *±* median absolute deviation. The significance of the difference between the distributions was assessed using Wilcoxon’s test with Bonferroni correction for multiple testing (n.s: non significant, P-value>0.05, *:0.01<P-value<0.05,**:0.001<P-value<0.01, ***: P-value<0.001). **e** Confocal images of TMRM fluorescence of each BJ5ta fibroblast cell line as indicated.

### MMUT-deficient cells show a mitochondrial, but no growth phenotype

In a high glucose, high glutamax (glutamine-alanine dipeptide) medium (DMEM mixed with M199, see Methods), we found no significant difference between MUTKO and WT growth rates (Fig. 2 **c**). However, tetramethylrho-damine methyl ester (TMRM) fluorescence, which accumulates in healthy mitochondria and measures mitochondrial potential^23^, indicated that BJ5ta-MUTKO cells had significantly (≥20%) lower mitochondrial potential than BJ5ta-WT fibroblasts (Fig. 2 **d**). Similarly, in confocal microscopy characterization, although mitochondria of MUTKO cells displayed a similar aspect ratio (ratio of the major and short axis of an ellipse fitted to an object) as controls cells, they were slightly more circular (measured by the form factor, the inverse of circularity; its minimum value is one for a perfect circle^24^; Fig. 2 **e** and Supplementary figure 2). Although MMUT-deficiency reduces mitochondrial potential, it does not induce a growth phenotype in standard medium.

### Changing medium conditions does not disrupt growth in MMUT-deficient cells

To induce growth disruption, we forced MMUT-deficient cells to rely more on respiratory metabolism, specifically by substituting glucose with galactose to increase reliance on oxidative phosphorylation^25–27^. This reduced the growth rates of BJ5ta-MUTKO cells to a similar extent as controls (Fig. 3 **a**), suggesting that the respiratory defect of MMUT-deficient cells is not severe.

**Figure 3.**
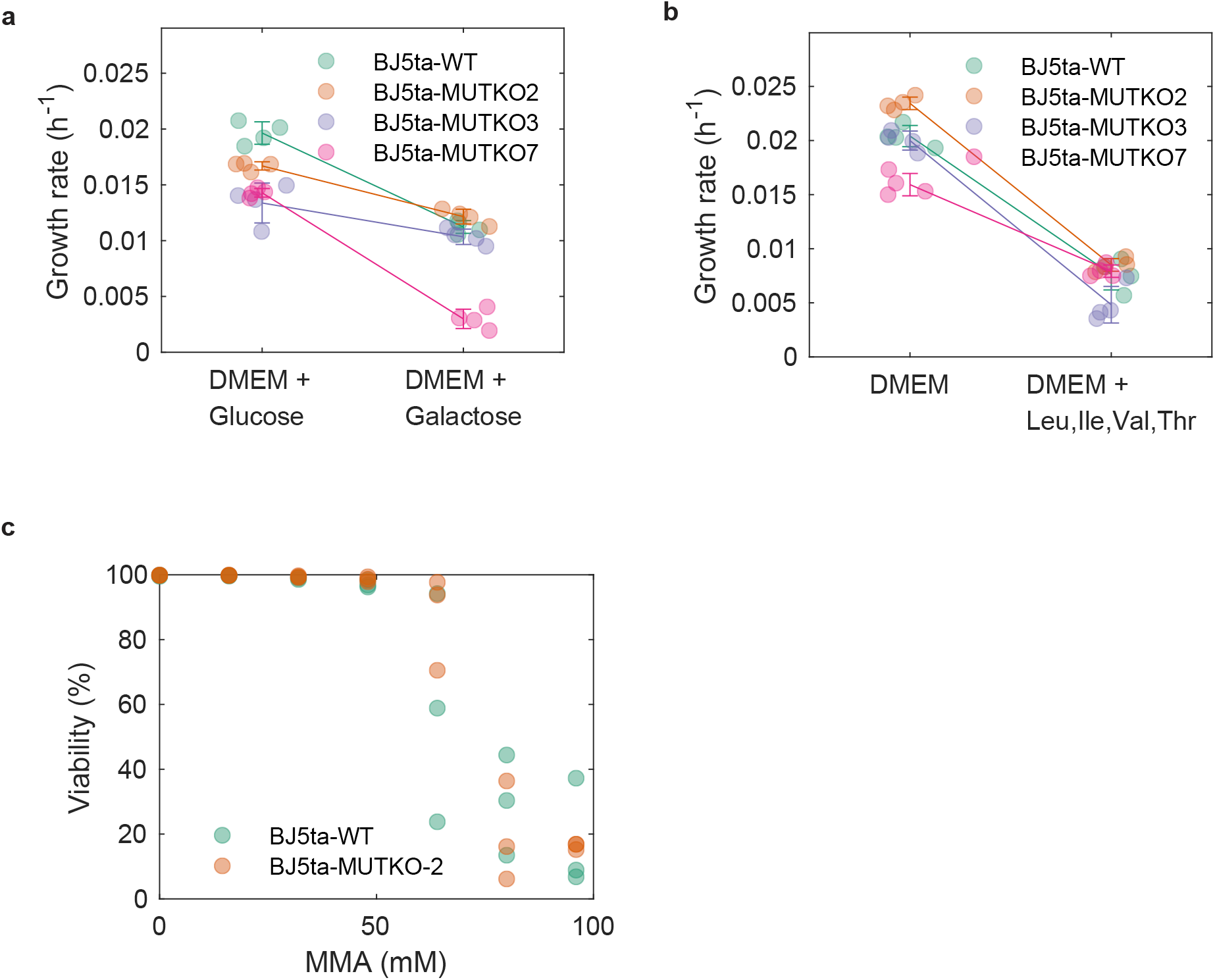
Growth and viability in non-standard media. **a** Growth rates of the BJ5tA fibroblasts were measured in normal DMEM with glucose, and when the carbon source was replaced with galactose. The circles are the raw data points, the error bars are the mean and standard deviations of the raw data. **b** Growth rates of the fibroblasts were measured in DMEM after adding leucine, isoleucine, valine and threonine each at a final concentration of 4 mM. **c** Viability as a function of the dose of methylmalonate (MMA). The viability was assessed after 72h and the experiment was performed in a pH controlled manner (i.e. the acidity was neutralised with NaOH).

Because dramatic response to addition of precursor amino acids (threonine, valine, isoleucine) were observed in MMUT-deficient mice^28^, we next attempted increasing these pools (and of leucine) in the medium. Supplementation with all four amino acids jointly (Fig. 3 **b**) and individually (Supplementary figure 3 **a** and Supplementary table 1) reduced the growth rates of all BJ5ta cell lines to a similar extent. Finally, to assess whether high concentrations of disease-related metabolites strongly affect viability in this cell type, we used medium with up to 100 mM MMA. However, we did not observe differences in viability between mutant and control cells (Fig. 3 **c** and Supplementary figure 3 **b**). Hence, in our cell models, MMUT deficiency does not affect growth in these previously tested conditions, and the elucidation of growth phenotypes requires quantitative modeling.

### Metabolic modeling predicts that MMUT-deficient cells compensate through anaplerotic pathways

We used structural sensitivity analysis^29^ to predict which fluxes change when MMUT is perturbed (see **Methods**). This identified 380 out of the 5957 reactions featured in the model to be changed significantly (Fig. 4 and Supplementary table 11). Specifically, for the *MMUT* deficient background, we predicted that: (i) fluxes in valine and threonine catabolism decrease, thereby increasing the extracellular concentrations of non-catabolizable pathway intermediates such as propionate and 2-methylcitrate; (ii) anaplerotic TCA cycle fluxes, such as the flux from glutamate to succinyl-CoA through alpha-ketoglutarate dehydrogenase, the pyruvate dehydrogenase flux, and the citrate synthase flux increase to compensate for the reduced MMUT flux; and (iii) extracellular concentrations of alpha-ketoglutarate and glutamate are reduced because of an increased intracellular need, for example, to feed in the TCA cycle. Such compensatory changes could explain the mild phenotypes observed in BJ5ta-MUTKO fibroblasts.

**Figure 4.**
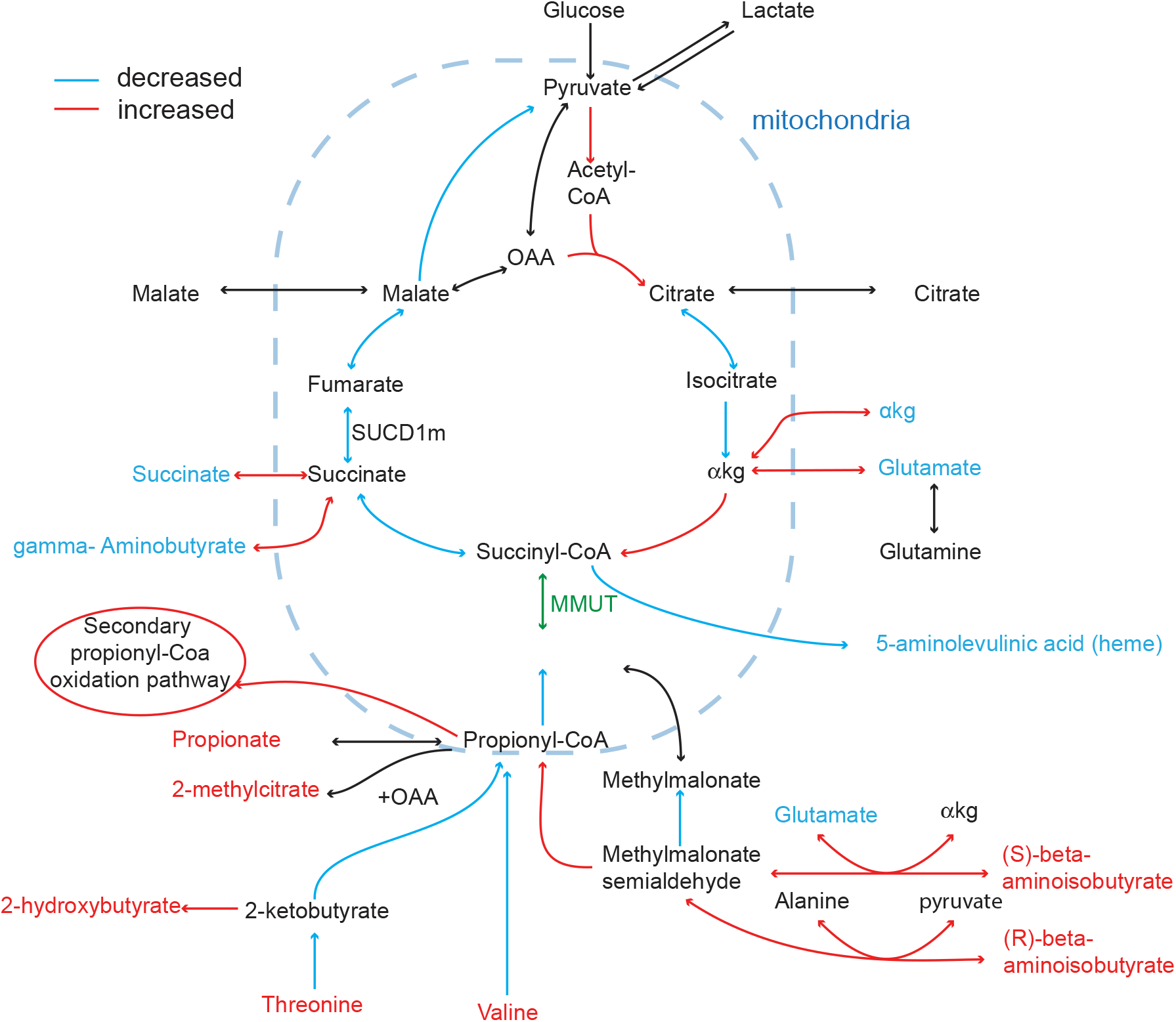
Prediction of the effect of MMUT deficiency on the rest of the metabolic network. Selected metabolic fluxes and extracellular metabolites predicted to be increased (red) or decreased (blue) upon perturbation of methylmalonyl-CoA mutase (MMUT) using sensitivity analysis. OAA: oxaloacetate; *α*kg: alpha-ketoglutarate.

### MMUT-deficient cells increase purine/pyrimidine and glutamine metabolism

To test the model predictions, we used a modified version of the tetrazolium-based redox assay, called Biolog phenotype microarrays^30^. These assays measure NADH production when cells grow on various metabolites; absorbance is proportional to the NADH reductase activity and the number of living cells. For most metabolites, we found an increased signal in BJ5ta-MUTKO fibroblasts, in particular for MUTKO2, compared to WT (Fig. 5 **a**). Lenth analysis of unreplicated factorials^31^ (see **Methods**) identified all metabolites that were significant for at least one cell line (Supplementary table 2 and Fig. 5 **a**).

**Figure 5.**
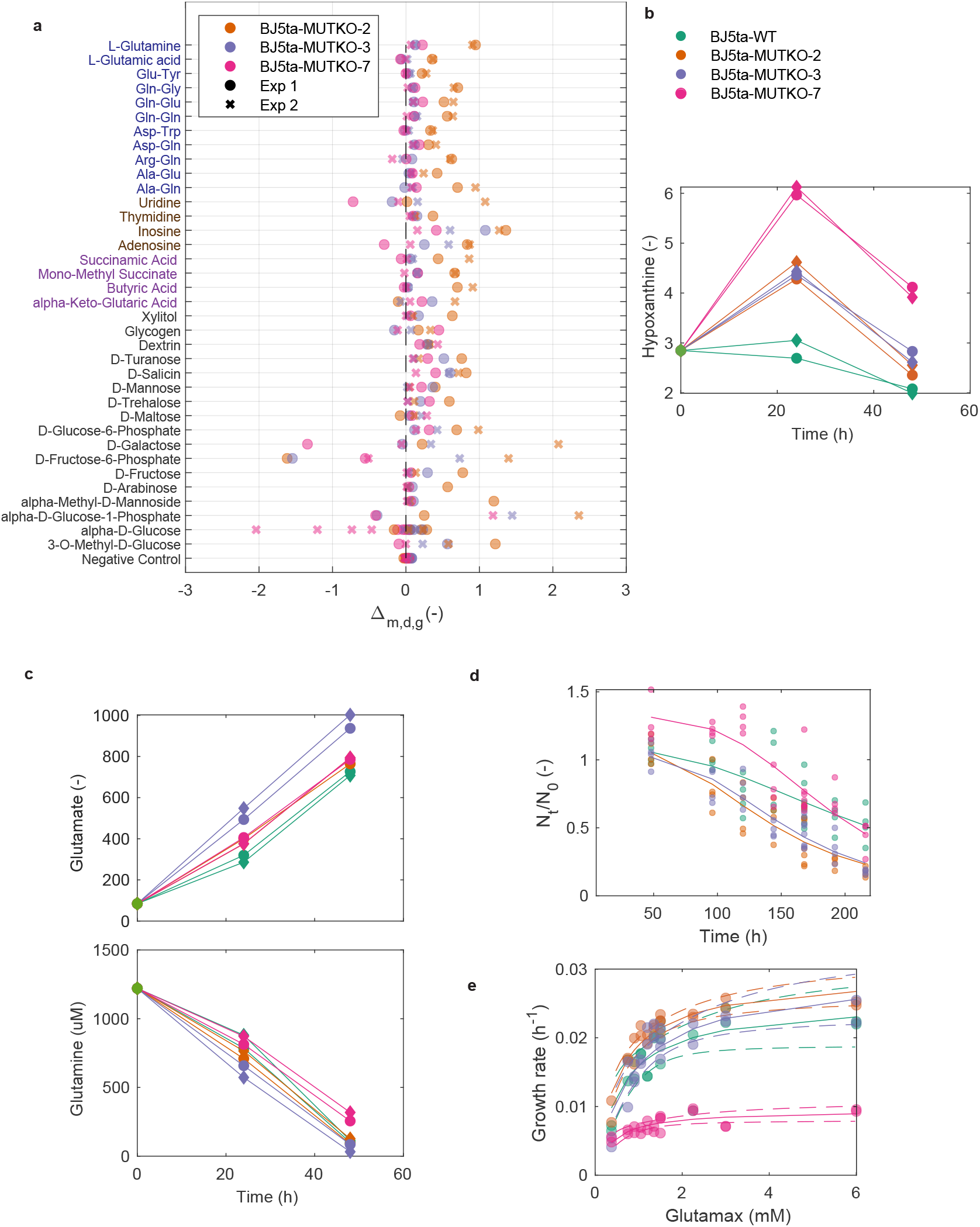
Metabolic dependencies of MMUT-deficient BJ5ta fibroblast cells. **a** Biolog phenotype microarrays for BJ5ta cells. Δ_*m,d,g*_ represents the difference in absorbance between mutant (g) and WT for each metabolite (m) on day (d). Absorbance was measured after 24h. Metabolites were selected by Lenth statistical analysis (see **Methods**). Results from two independent experiments (symbols). **b** Hypoxanthine secretion measured in the spent medium of the four cell lines indicated. **c** Glutamate secretion (top) and glutamine consumption (bottom) detected in the spent medium of BJ5ta fibroblast cells; symbols as in **b. d** Fraction of viable cells in a medium containing neither glutamax nor glutamine as a function of time. Dots: well; solid line: fit of a 3-parameter logistic curve. For parameter estimates and associated confidence intervals, see Supplementary table 9. Symbols as in **b. e** Growth rate as a function of glutamax concentration. Solid line: fit of a Michaelis-Menten curve. Estimated parameters and associated confidence intervals can be found in Supplementary table 4. Symbols as in **b**.

Only inosine, D-salicin and alpha-D-glucose-1-phosphate were common to all MUTKO cell lines, with inosine and D-salicin showing the biggest increase. Overall, however, upregulated metabolites cluster in four categories: L-glutamine, glutamate and related dipeptides (in MUTKO2 and MUTKO7); purines and pyrimidines (adenosine, which is one step away from inosine, and to a lesser extent uridine and thymidine); organic acids (alpha-ketoglutarate, succinamic acid, butyrate and mono-methyl succinate, mostly in MUTKO2); and sugars.

We performed the same experiment on CRISPR-Cas9 modified HEK293 cells deficient in MMUT, MMAB (an enzyme involved in MMUT cofactor synthesis), or both. We observed similar trends as in fibroblasts, in particular, upregulation with glutamine dipeptides (Gln-Gln, Gln-Gly) and adenosine (Supplementary figure 4).

Our modeling and Biolog results suggested that MUTKO cell lines have (i) an increased inosine metabolism and (ii) an increased L-glutamine/glutamate metabolism. To pursue these hypotheses, we measured secretion and uptake rates in the spent medium of BJ5ta cell lines. Secretion of hypoxanthine (derived from inosine) was increased for MUTKO cell lines compared to WT at 24h, followed by a decay (Fig. 5 **b**). Compared to WT, all MUTKO cell lines significantly increased glutamate secretion (Fig. 5 **c**, Supplementary table 5 and Supplementary table 6), and MUTKO2/3 cells also significantly increased glutamine consumption (Supplementary table 7 and Supplementary table 8).

We hypothesized that MUTKO cells depend more on glutamine to compensate for the reduced flux through the TCA cycle. Indeed, MUTKO2 and MUTKO3 died faster in a glutamine-free medium, while growth rates for increasing doses of glutamax were not significantly different from WT (Fig. 5 **d**,**e**, Supplementary table 9, and Supplementary table 4). In contrast, MUTKO7 grows well in the absence of glutamine at early time points (Fig. 5 **d**) and its growth rate is not strongly stimulated in increasing concentrations of glutamax (Fig. 5 **e**), suggesting it does not rely on glutamine for growth. Overall, the results are consistent across experiments per cell line, but they illustrate clonal variability resulting from modifying cells with CRISPR-Cas9.

### *In silico* prediction of MMUT function

To interpret these results, we analyzed under which conditions the reversible reaction catalyzed by MMUT (MMMm) transforms methylmalonyl-CoA into succinyl-CoA (positive flux in the model), thereby fulfilling its anaplerotic function. Flux variability analysis^32^ predicted a positive flux through MMMm either with ATP or biomass flux at their maximum, or a linear combination of both (Fig. 6 **a** and Supplementary figure 5 **a**). Also, for limiting uptake of glucose and glutamine from the medium, maximal growth of MUTKO cells is predicted to decrease compared to WT, but the reduction to 99% of WT is small (Fig. 6 **b**). Importantly, this differential growth arises when the MMMm flux becomes positive (Supplementary figure 5 **b**). This is consistent with a decreasing maximum growth rate when there is an increasing fixed positive flux through MMMm (Fig. 6 **c**). To explain the upper limit to the MMMm flux for maximal growth, we identified which components of the biomass reaction limit growth (see **Methods**). We predicted growth with and without each component *i*, yielding a positive difference of growth rates 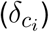 when compound *i* limits growth. As the MMMm flux increases, 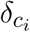 for direct (isoleucine, valine and threonine) and more distant (dCTP, dTTP, CTP, UTP) precursors of MMMm increases (Supplementary figure 6). Hence, a high flux through MMMm may be disadvantageous for the cell, explaining why MMMm will function in the forward direction only under stringent conditions.

**Figure 6.**
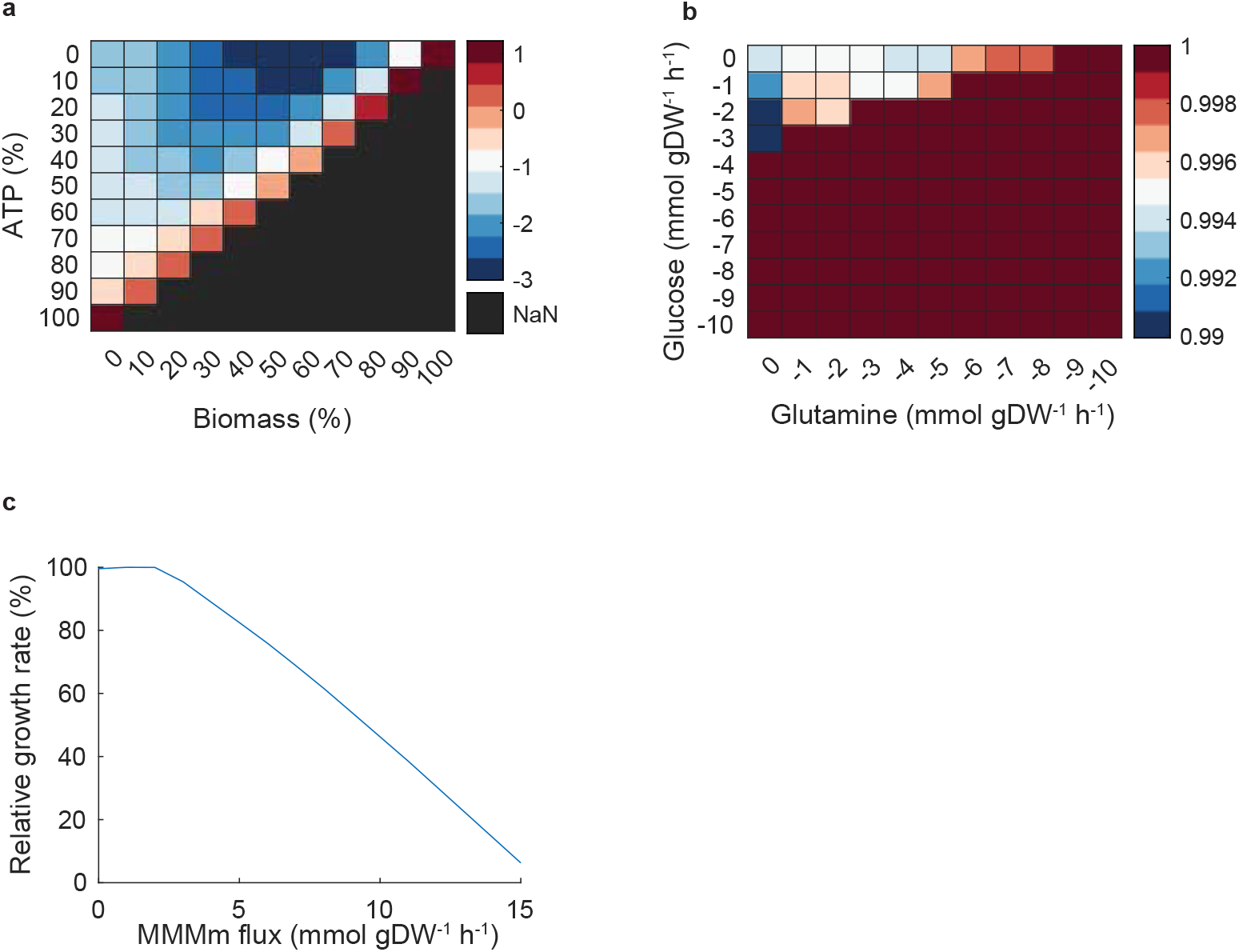
*In silico* prediction of MMMm flux in different conditions. **a** Flux variability analysis of the MMMm reaction as a function of the minimum ATP and biomass production required. Average of the minimum and maximum flux value through the MMMm reaction. **b** Flux balance analysis predictions for the maximum growth ratio of MMUTKO vs WT as a function of glucose and glutamine exchanges; negative values indicate uptake. **c** Percentage of maximum growth rate as a function of fixed MMMm flux.

### Genetic interactions with *MMUT*

Finally, to investigate genetic interactions, we predicted single-reaction deletions that decrease the growth ratio (see **Methods**). We found only 13 out of the *¥*5’400 model reactions to interact with the MUTKO phenotype (Fig. 7 **a**). They mostly relate to the secondary oxidation of propionyl-CoA and to respiration (respiratory chain complexes and oxygen transport), resulting in the smallest growth ratio of 88%. Because there can be many-to-many relations between genes and reactions, we repeated the analysis by knocking out individual genes, yielding consistent results (Supplementary table 12).

**Figure 7.**
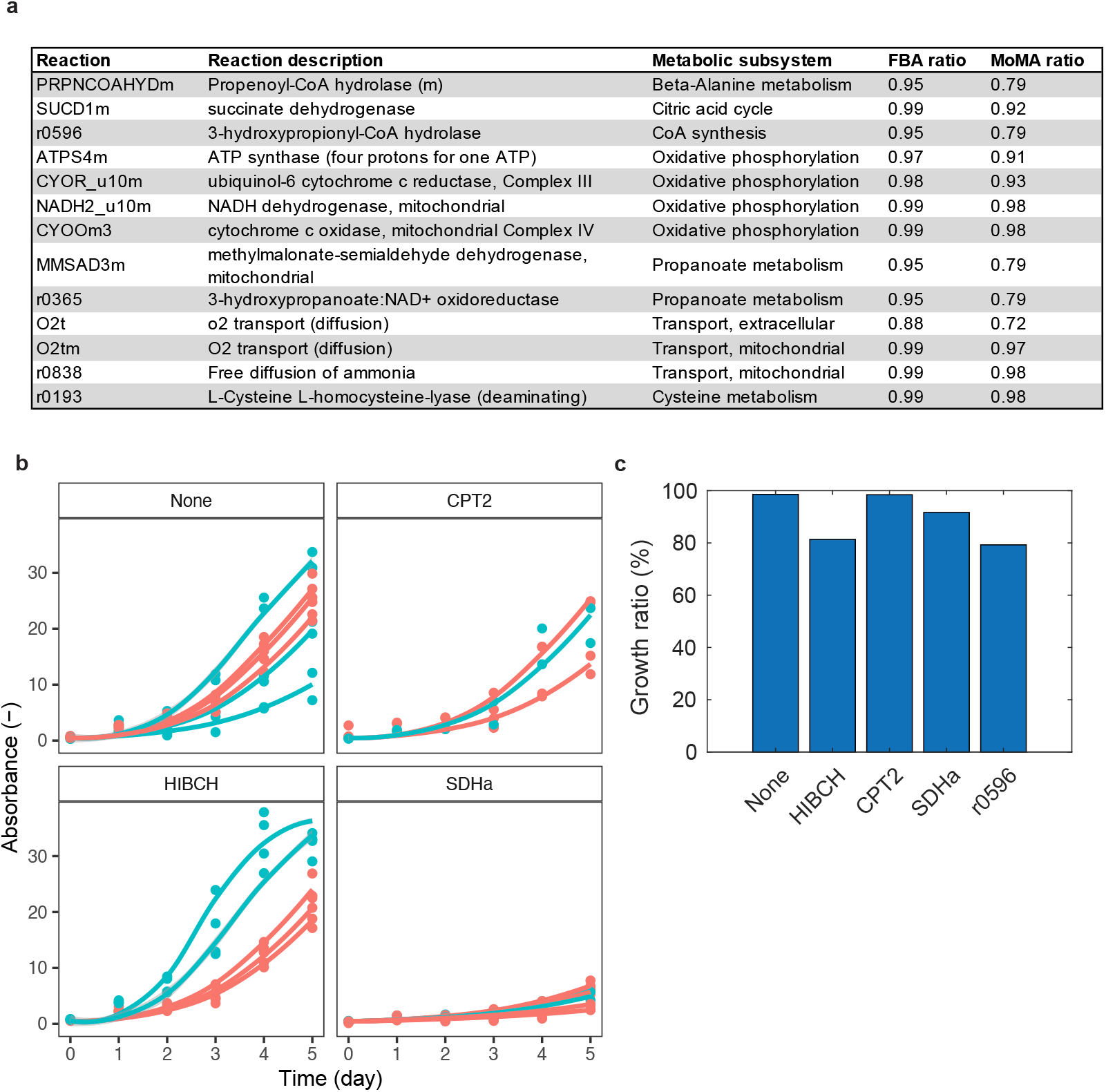
Genetic interactions with *MMUT*. **a** Single-reaction deletion *in silico* experiment. Reactions that create a growth defect in mutant vs WT simulated in a DMEM medium and corresponding growth ratio (see **Methods**). **b** Growth of WT (blue) and *MMUT* - deficient (red) HEK293 cells when an additional gene is knocked-out: carnitine palmitoyltransferase II (*CPT2*), 3-hydroxyisobutyryl-CoA hydrolase (*HIBCH*), succinate dehydrogenase complex flavoprotein subunit A (*SDHa*) or when no other gene is knocked-out (None). Cell growth was measured using the Crystal Violet assay; absorbance is proportional to the number of cells. Dots: wells; curve: fit of a non-linear mixed effect model based on a logistic curve (see **Methods**). **c** MoMA predictions of growth ratio of *MMUT* -deficient cells to WT when another gene is knocked-out (horizontal axis) or when the flux through reaction r0596 is blocked.

Based on the predictions, we created CRISPR-Cas knock-outs of *HIBCH, CPT2* and *SDHa* as they belong to very different metabolic pathways in the WT and MUTKO background of HEK293 cells. To control for potential side effects of the CRISPR procedure we used the parental cell lines as well as control cells that went through one unsuccessful round of CRISPR-Cas9 (see **Methods**).

Growth rates of each cell line were measured (Fig. 7 **b**) and we assessed the significance of genetic interactions between MMUT and CRISPR-Cas knockouts with a non-linear mixed effect model for cell growth measured by the Crystal Violet assay (see **Methods** and Supplementary table 13). We found a significant interaction between *HIBCH* and *MMUT* (Supplementary table 14). Importantly, the HIBCH enzyme catalyzes multiple reactions: the hydrolysis of 3-hydroisobutyryl-CoA in valine catabolism, as well as the hydrolysis of 3-hydroxypropionyl-CoA (reaction r0596) in the secondary propionic acid oxidation pathway. Looking back in the model, we found the growth difference to be explained by reaction r0596 (Fig. 7), suggesting that the secondary oxidation pathway of propionyl-CoA interacts with loss of MMUT function, consistent with our previous results.

## Discussion

The exact pathomechanisms behind the adverse effects of MMAuria remain largely unknown, also because of the interconnected nature of the human metabolic network. Our approach is distinct from earlier studies of individual enzymes^10–12^ in analyzing the network effects of perturbations with cellular and computational models. In addition, we focused on simple observables (growth and viability) that integrate multiple signals in the cell. Assays specifically measuring mitochondrial function could be alternatives for future studies because reduced absorbance of MTT (a tetrazolium-based assay similar to the Biolog assay) has been found in MUT-KO HEK293 cells in a propionate enriched medium^33^.

We used skin fibroblasts as the basic cell type to study MMAuria although they are not primarily affected by the disease. However, patient fibroblasts are commonly studied in this context. For example, fibroblasts from patients with respiratory chain defects showed a phenotype in galactose medium, despite not being the primarily affected cell type^34^. Specifically, MMUT is expressed ubiquitously^35^ and the pathway is functional in fibroblasts^36^. In addition to them being an easy cell type to handle experimentally, candidate results could be tested on patient fibroblasts in future studies.

Our BJ5ta-MUTKO fibroblasts showed mitochondrial impairment as expected from clinical observations^37^. However, none of the standard treatments to elicit differential growth defects in MMUT-deficient cells (galactose, amino acid precursors, and MMA) yielded a growth phenotype. This suggests more complicated mechanisms than the “toxic metabolite” hypothesis alone. Note, however, that our compound list was not exhaustive. For instance, 2-methylcitrate had inhibitory effects of enzymes in the TCA cycle in some studies^19,20^.

In the Biolog assay, NADH metabolism was generally increased in BJ5ta-MUTKO fibroblasts and HEK293 cells, with the biggest effects for purine/pyrimidine-based metabolites and glutamate/glutamine. For purine metabolites (inosine and adenosine), hypoxanthine secretion was increased in the spent medium of BJ5ta-MUTKO cells. Hypoxanthine could be the unnecessary byproduct of the transformation of inosine into hypoxanthine and ribose-1-phosphate because MUTKO-cells would have an increased need for ribose-1-phosphate, part of the pentose phosphate pathway. Purine metabolites were already associated to mitochondrial deficiency since inosine and uridine were changed in the spent medium of myotubes treated with respiratory chain inhibitors^38^. Hence, this part of metabolism could deserve further investigation.

Besides increased NADH metabolism of glutamine related dipeptides, the viability of two of the BJ5ta-MUTKO fibroblasts (MUTKO2 and MUTKO3) decreased faster in a glutamine-free medium, a stringent condition for the cells. However, we observed no growth difference between these two BJ5ta-MUTKO fibroblasts and WT for various doses of the glutamine-alanine dipeptide glutamax, even if glutamine consumption was increased in BJ5ta-MUTKO fibroblasts. Consistently, we predicted a 1% difference in growth between MUTKO and WT could be obtained by reducing amounts of glutamine in the medium. Consequently, BJ5ta-MUTKO fibroblasts might compensate for MMUT deficiency by increasing their reliance on glutamine metabolism, but with an effect on viability only when cells are stressed. Similarly, a reduced survival in mutant flies was reported upon starvation in a CRISPR-Cas9 Drosophila model of succinyl-CoA synthetase deficiency, an enzyme in the TCA cycle^39^. This illustrates how the medium can be crucial when studying a metabolic disease.

Regarding glutamate metabolism, we observed increased glutamate secretion in BJ5ta-MUTKO fibroblasts. In liver cancer cells, which also display increased reliance on glutamine metabolism, increased glutamate secretion was hypothesized to be a consequence of increased flux through the *de novo* purine synthesis pathway, which also transforms glutamine into glutamate besides producing purines^40^. This hypothesis would be consistent with our findings.

To interpret these results, we used constraint-based modeling. We predicted increased anaplerotic fluxes towards the TCA cycle and reduced oxidative flux in the TCA cycle. This is consistent with glutamate dehydrogenases being upregulated in the hepatic proteome of MMAuria patients^41^ and in MMUT-deficient fibroblasts^42^. Interestingly, we predict a global reduced oxidative TCA cycle flux consistent with a recent study^42^, except for alpha-ketoglutarate dehydrogenase for which we predict increased flux, while the protein amount was decreased in MMUT-deficient primary fibroblasts^42^. This discrepancy could due to protein-protein interactions between OGDH and MMUT^42^ or kinetics that are not captured by the model.

In predicting which metabolic reactions would interact with MMUT to create a growth phenotype, we found that reactions involved in oxidative phosphorylation and the secondary propionyl-CoA oxidation pathway interact with MMUT deficiency. In one experimental test of the predictions, HIBCH, part of the secondary-propionyl-CoA secondary oxidation pathway, indeed interacted with MMUT by diverting flux to this secondary pathway. This pathway was transcriptionally activated in *C. elegans* on vitamin B12 deficient diets^43^. This illustrates how the combination of modeling and experiments yields interpretable hypotheses.

Finally, we observed clonal variability of the BJ5ta-MUTKO cells that was not apparently caused by genetic modifications due to CRISPR-Cas9 off-target effects. It could be the result of the clonal selection of single cells. This variability complicated the interpretation of our experiments, but evidenced the need to test multiple mutants to avoid over-interpreting results. Although we cannot exclude that the effects we report are due to the CRISPR-Cas9 modification and not the MMUT deficiency, the findings are consistent with other models of the disease, as discussed above.

Overall, MMUT-type methylmalonic aciduria has complex and mild effects in fibroblasts, which could be linked to the reduced TCA cycle flux. This work suggests novel candidate metabolites and pathways to test in the future.

## Methods

### Cells

The human foreskin fibroblasts immortalized with hTERT were bought from ATCC (BJ5ta, CRL 4001) and were cultured as recommended in medium A : DMEM with Glutamax (31966047, Thermofisher), mixed in a 4:1 ratio with M199 (Thermofisher, 41150020), 10% (v/v) Fetal Bovine Serum (FBS) and 0.01 mg/ml hygromycin B. Human Embryonic Kidney (HEK293) cells were bought from ATCC (HEK293 CRL 3216) and cultured in DMEM with Glutamax, 10% Fetal Bovine Serum and 1X Antibiotic Antimycotic (15240062, Thermofisher). Cells were kept at 37°C under 5% CO2 atmosphere in 100 mm dishes. The cells were tested negative for mycoplasma contamination using Mycoplasmacheck service (Eurofins).

### CRISPR-Cas9 generation of knockouts

To generate knock-out (KO) in BJ5ta and HEK293 cells, we used CRISPR/Cas9 technology^44^. BJ5ta cells and HEK293 cells were transfected using the 4D nucleofector (Lonza) and Neon transfection system (Thermo Fisher Scientific), respectively. Single guide RNAs (sgRNAs) were provided either as gBlocks (IDT technologies)^45^ with a Cas9 plasmid (Addgene: 62988) or directly as a ribonucleoprotein complex (Cas9 Nuclease: A36496; IVT gRNA: A29377). After 48 hours, cells were transferred to a 96-well plate to obtain an average of 1 cell/well and cultured for clonal expansion. Sanger sequencing was performed for selection on genomic DNA. SgRNAs targeting MMUT in BJ5ta cells were designed to target Exon 3 of *MMUT* : ATAGTAACTGGAGAAGAACA (sgRNA1) and ACGATGTGTCGCCAGATCAA (sgRNA 2). In HEK293 cells, the sgRNA targeting MMUT was ATTCCTTTAGTATATCATTTTGG. Knock-out of HIBCH, CPT2 and SDHa was performed both on wild-type (WT) HEK293 cells and on successful MUT-KO HEK293 pre-generated cells. Guide RNAs were designed as follow: HIBCH: GCAGATTTATCCACAGCTAAAGG, CPT2: TGTTGGTTGC-CCGGGTGAGCTGG, SDHa: ACGTCTGCCCACACCAGCACTGG. Control cells were WT HEK293 or MUT-KO HEK293 cells which have gone through CRISPR/Cas9 editing but did not show any knockout of our targeted genes.

### Validation of knockout using Western blotting

Lysates from a confluent T75 were obtained using RIPA buffer and were mixed with RIPA lysis buffer and 2X Lammli buffer (containing 5% (v/v) *—*-mercaptoethanol) to obtain a protein concentration of 1.25 µg/µl. Electrophoresis was performed on a SDS page gel (10% polyacrylamide) in Tris-glycine buffer using 20 µl of protein. Proteins were transferred using the semi-dry method onto a nitrocellulose membrane (Whatman, GE Healthcare). The membrane was blocked in the blocking buffer (5% skimmed milk, 20mM Tris base, 150 mM NaCl, 0.2% Tween 20) for one hour at RT, incubated with primary antibodies (anti-MMUT: ab67869, Abcam/ anti-ACTIN, Sigma, A1978) overnight at 4°C diluted 1:500 in blocking buffer and incubated with secondary antibodies (anti-mouse HRP, Sc516102-CM, Santa Cruz) diluted 1:5000 in blocking buffer for one hour at RT. Signal was detected using the Clarity ECL Substrate (Biorad, 1705060S) and ChemiDoc™ Touch Imaging System.

### Whole-genome sequencing and analysis

DNA was extracted using a blood and cell culture DNA mini kit (13323, Qiagen) from low passage cell lines, as recommended. For the sequencing, library preparation was performed using a KAPA HyperPrep Kit PCR-free (Roche) with a target insert size of 500 bp. Paired-end libraries were sequenced using a GFB Novaseq 6000 sequencer (Illumina, RTA Version: v3.4.4). Base calls were converted into FASTQ files using bcl2fastq v2.20.0.422 and further analysed to find the germline variants.

Raw reads are quality controlled. Sequencing adapters are removed from the reads and trimmed reads are aligned to the reference genome (GRCh38). Aligned reads are post-processed (removal of secondary alignments and PCR duplicates) and recalibration of quality scores is performed. Germline variant calling and filtering is performed for each cell line. Three separate variant callers are combined: Strelka, v2.9.2, GATK HaplotypeCaller, v4.1.6.0 and Varscan v2.4.3. Only calls which are detected by at least 2 out of 3 tools are kept. Variant annotation is performed by VEP (v103.1). Only variants with a frequency higher than 30% are used since we expect 50% and 100% variant frequency for germline variants. Variants are compared across cell lines and only variants for which not a single read can be found in the WT cell line are considered novel in the mutant cell line.

### MMUT activity assay

MMUT enzyme activity assay was performed using crude cell lysates as described^42^.

### Cell proliferation and viability experiment in different media

These experiments were performed in DMEM without glucose, glutamine and phenol red (Thermofisher, A143001). Unless otherwise stated, this medium was supplemented with 25 mM glucose (Sigma, G5388), 1mM Glutamax (thermofisher, 35050061), 0.04 mM phenol red (ATCC, PCS-999-001) and 10% (v/v) dialyzed FBS (Thermofisher, 26400044).

2500 cells were seeded per well of a 96-well plate (IBIDI, 89626) in medium A. The following day, the wells were rinsed and replaced with the desired medium. Cell proliferation was assessed by imaging cells using microscopy (Supplementary figure 7 **a**) The cells were imaged using a Nikon Ti2 microscope and a Andor Sona CSC-002200 camera at +150 µm above the focal plane using bright field imaging and using a magnification of 6x (4x plan Apo *λ* objective *na* = 0.2, and 1.5x zoom). Three or four days later, the cells nuclei were stained using the Nucblue stain (Thermofisher, R37605) as recommended and imaged. Cells were counted by segmenting the bright-field and fluorescent images using Fogbank algorithm^46^ (segmentation parameters: Supplementary table 15). The same number of cells are counted using bright-field imaging and nuclear staining when the number of cells in the field-of-view is below 1500 cells (Supplementary figure 7 **b**) at the beginning of the experiment.

To assess the viability of the cells upon treatment with a compound or a nutrient, cells were stained with a membrane impermeable dye Sytox (Thermofisher, S7020, diluted 1:5000), on top of the membrane permeable dye (Nucblue). A cell is then considered viable if it stained by the membrane permeant dye but not by the membrane impermeant dye.

### TMRM experiment and confocal imaging

Cells were seeded in a 96 well-plate. One day later, cells were stained with 50 nM Tetramethylrhodamine, Methyl Ester, Perchlorate (TMRM) (Thermofisher, T668) diluted in medium A devoid of phenol red for 30 minutes at 37°C in the dark. After rinsing, fluorescent (20x Plan Fluor objective, excitation : 531 nm, Cy3 light filter) and bright-field images were acquired. A mask of individual cells was obtained by segmenting the bright-field image. The total fluorescence of a cell was computed by summing the blanked fluorescence of each pixel inside the cell, where the blank is the fluorescence of the background.

Mitochondria were also imaged using a Leica confocal microscope, using a HC PL APO 63x/1.40 oil objective. The difference in their shapes were assessed using descriptors such as the form factor and the aspect ratio of the mitochondria, which were computed using an existing script ^24^.

### Biolog phenotype microarray and Lenth statistical analysis

20000 cells per well were resuspended in the assay medium (IF-M1(Biolog Inc.,BL-72301), 0.3 mM L-glutamine, 5% FBS) and seeded in Biolog plates PMM1(BL-13101) and PMM2 (BL-13102). After 46 hours of incubation at 37°C, 5% CO2, the dye MB (Biolog Inc,BL-74352) was added. After 24h, the absorbance of the plate was measured at 590 nm and 750 nm. *a*_*m,d,g*_ denotes the absorbance at 590 nm subtracted with the absorbance at 750 nm, where *m* is the metabolite considered, *g* is the cell line and *d* is the day the experiment.

Lenth analysis^31^ of unreplicated factorials was performed on data adjusted for systematic differences between days, metabolites and baseline abundance in wild-type:

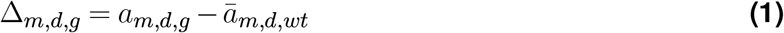

where, *ā*_*m,d,wt*_ is the average absorbance of metabolite *m* in wild-type on day *d*. Candidate metabolites fulfill |Δ_*m,d,g*_| *> SME*, where *SME* is the simultaneous margin of error and is computed:

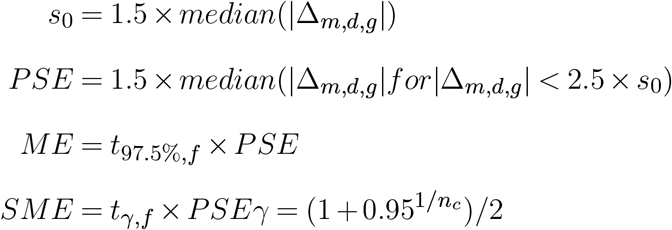

where *s*0, *PSE*, and *ME* are the standard deviation, pseudo standard error and margin of error, respectively. *f* is the number of degrees of freedom, which is *f* = *n*_*c*_*/*3, where *n*_*c*_ is the number of contrasts.

### Metabolite detection experiment and parameter estimation

10^6^ cells were seeded per 100 mm dish in duplicates. One day later, dishes were rinsed and 11 mL of a medium containing a DMEM free of glucose and glutamine, with 6.25 mM glucose, 1 mM glutamine, 50 uM inosine, 10% dialysed FBS and hygromycin B. In each dish, two times 1mL of spent medium samples were taken at 24 and 48 hours. Immediately after retrieval, samples were counted and were flash-frozen in liquid nitrogen and stored at -80°C for further analysis.

Hypoxanthine concentration was measured using the inosine fluorometric assay kit (Abcam, ab126286) without the converter enzyme. Glutamine and glutamate were assayed using a glutamine assay kit (Abcam, ab197011), as recommended for glutamine and without the hydrolysis enzyme mix for glutamate.

For glutamate and glutamine, we estimated the secretion and consumption rate, respectively. For glutamate:

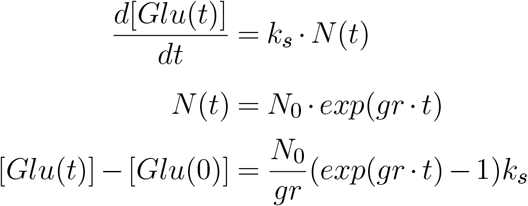

For glutamine:

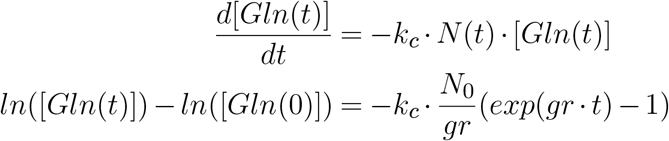

[*Gln*(*t*)] and [*Glu*(*t*)] are the concentration of glutamine and glutamate in the spent medium respectively, *k*_*s*_ and *k*_*c*_ are the secretion and consumption rate per cell, *N* (*t*) and *N*_0_ are the number of cells at time *t* and initial time respectively, *gr* is the growth rate. We used standard linear regression to estimate the parameters. For glutamine, only the sample at 24 h was selected since at 48 h, there was no glutamine left in the medium.

### Crystal violet assay

12500 cells were seeded on Poly-L-Lysine well of a 24 well plate in duplicates. Cells were counted using the crystal violet assay (Sigma, C6158) at day 0,1,2,3,4 and 5. For each measurement, the well was washed using DPBS, fixed in 200 µL of 4% PFA for 10 minutes, incubated with crystal violet (0.2 %) for 20 minutes. The plate was sinked three times under water to remove the extra crystal violet. 1000 µL of 1% SDS was then added to each well followed by 30 minutes of shaking at RT. Absorbance was measured at 540 nm in a Tecan microplate reader.

A non-linear mixed effect model was then fitted to the data. For a cell line *i*, the following non-linear mixed effect model was estimated:

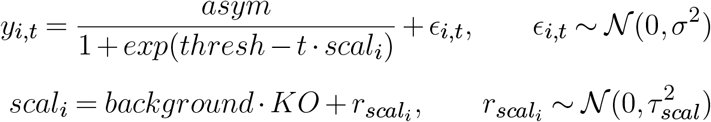

where *background* and *KO* describe whether the cell line is *MMUT* deficient and if another gene was knocked-out respectively.

### *In silico* modeling

We performed constraint-based modeling with human reconstruction Recon 2.2^47^ and simulated DMEM medium (see Supplementary table 16). To predict maximum growth and maximum ATP generation, we used standard flux balance analysis^48^. Varying amounts of glucose and glutamine were modeled by fixing the uptake fluxes as indicated. Reaction deletions were modeled by fixing the respective fluxes to zero.

To understand which compounds limit growth, we added artificial uptake reactions for components *C*_*i*_ with upper bound fixed to 1. If only *C*_*i*_ limits growth, biomass flux increases by 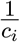, where *c*_*i*_ is *C*_*i*_’s stoichiometric coefficient in the biomass reaction. We computed the resulting biomass increase as 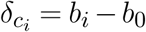, where *b*_*i*_ (*b*_0_) is the biomass flux after (before) adding *C*_*i*_.

To predict the spreading of the MMUT perturbation in the network, we used structural sensitivity analysis^29^. Specifically, we computed the adjustments *d*_*k,e*_ of exchange fluxes *e* to a perturbation ***δ***_**k**_ of the MMMm reaction **k**, yielding sensitivities

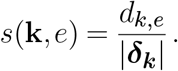

To decide whether these sensitivities are significant, we compared them to the distribution of the sensitivities to all internal reactions *i* being perturbed *s*(*i, e*). With the median *med*_*e*_ and median absolute deviation (MAD) *mad*_*e*_ of this distribution, we classified metabolite *e* as a biomarker when

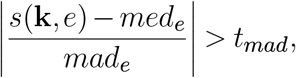

with *t*_*mad*_ = 30, similar to previous methods for outlier detection^49^.

## ACKNOWLEDGEMENTS

We thank Fabian Rudolf for initial discussions about the project, Hans-Michael Kaltenbach for the advice and for performing the statistical analyses and Lukas Stoob for the methylmalonyl-CoA mutase activity assay. DSF received financial support from the Swiss National Science Foundation (310030-200798) and the University Research Priority Program of the University of Zurich (URPP) ITINERARE – Innovative Therapies in Rare Diseases.

## AUTHOR CONTRIBUTIONS

DSF and JS conceived of and supervised the work. CR performed the experiments and did the modeling. CR and CB supported CRISPR-Cas9 mediated creation of BJ5ta MMUT deficient cell lines, CB and TF generated the HEK293 CRISPR-Cas9 cells. CR, CB and FT performed the crystal violet assays. The Lenth statistical analysis was performed by Hans Michael Kaltenbach. CR wrote the manuscript with support from DSF and JS.

## DATA SHARING POLICY

All relevant data has been provided in the manuscript as main or supplementary information. Any further supporting data can be supplied upon reasonable request to the authors.

