## Supplementary information for "Cellular and computational models reveal environmental and genetic interactions in *MMUT*-type methylmalonic aciduria"

### Supplementary tables

**Supplementary table 1. Parameter estimates**. IC50 parameter estimate and associated 95% confidence intervals.

| Amino acid | Cell lines | $log_2(IC50)$ | lower CI | upper CI |
|---|---|---|---|---|
| Valine | BJ5ta-WT | 5.02 | 4.46 | 5.59 |
| | BJ5ta-MUTKO2 | 4.85 | 4.31 | 5.39 |
| | BJ5ta-MUTKO7 | 4.98 | 4.36 | 5.61 |
| Threonine | BJ5ta-WT | 3.37 | 2.83 | 3.91 |
| | BJ5ta-MUTKO2 | 3.59 | 3.25 | 3.93 |
| | BJ5ta-MUTKO7 | 4.38 | 3.84 | 4.93 |
| Leucine | BJ5ta-WT | 3.22 | 2.83 | 3.61 |
| | BJ5ta-MUTKO2 | 2.97 | 2.67 | 3.27 |
| | BJ5ta-MUTKO7 | 3.27 | 3.02 | 3.53 |
| Isoleucine | BJ5ta-WT | 3.71 | 3.47 | 3.95 |
| | BJ5ta-MUTKO2 | 3.64 | 3.36 | 3.93 |
| | BJ5ta-MUTKO7 | 3.89 | 3.46 | 4.33 |

**Supplementary table 2. Candidate metabolites per BJ5ta cell line**. Derived using Lenth statistical analysis of unreplicated factorials using the Biolog phenotype microarray data.

| Cell line | Metabolite |
|---|---|
| BJ5ta-MUTKO2 | D-Glucose-6-Phosphate |
| | alpha-D-Glucose-1-Phosphate |
| | D-Galactose |
| | D-Arabinose |
| | Dextrin |
| | D-Mannose |
| | Xylitol |
| | Glycogen |
| | alpha-Methyl-D-Mannoside |
| | alpha-Keto-Glutaric Acid |
| | 3-O-Methyl-D-Glucose |
| | D-Fructose |
| | Succinamic Acid |
| | Butyric Acid |
| | Thymidine |
| | Mono-Methyl Succinate |
| | D-Trehalose |
| | D-Salicin |
| | Uridine |
| | Adenosine |
| | D-Turanose |
| | Inosine |
| | Asp-Gln |
| | L-Glutamic acid |
| | Glu-Tyr |
| | Ala-Glu |
| | L-Glutamine |
| | Ala-Gln |
| | Gln-Glu |
| | Gln-Gln |
| | Asp-Trp |
| | Gln-Gly |
| | Arg-Gln |
| BJ5ta-MUTKO3 | alpha-D-Glucose-1-Phosphate |
| | D-Fructose-6-Phosphate |
| | 3-O-Methyl-D-Glucose |
| | D-Salicin |
| | Adenosine |
| | Inosine |
| BJ5ta-MUTKO7 | D-Glucose-6-Phosphate |
| | alpha-D-Glucose-1-Phosphate |
| | D-Galactose |
| | alpha-D-Glucose |
| | Dextrin |
| | D-Fructose-6-Phosphate |
| | D-Maltose |
| | D-Salicin |
| | Uridine |
| | D-Turanose |
| | Inosine |

**Supplementary table 3. Parameter estimates**. Km parameter estimate for Glutamax growth rate dose response and associated 95% confidence intervals.

| Cell lines | $Km$ | lower CI | upper CI |
|---|---|---|---|
| BJ5ta-WT | 0.59 | 0.18 | 1.01 |
| BJ5ta-MUTKO2 | 0.54 | 0.38 | 0.70 |
| BJ5ta-MUTKO3 | 0.83 | 0.44 | 1.22 |
| BJ5ta-MUTKO7 | 0.37 | 0.16 | 0.58 |

**Supplementary table 4. Parameter estimates**. Km parameter estimate for Glutamax growth rate dose response and associated 95% confidence intervals.

| Cell lines | $Km$ | lower CI | upper CI |
|---|---|---|---|
| BJ5ta-WT | 0.59 | 0.18 | 1.01 |
| BJ5ta-MUTKO2 | 0.54 | 0.38 | 0.70 |
| BJ5ta-MUTKO3 | 0.83 | 0.44 | 1.22 |
| BJ5ta-MUTKO7 | 0.37 | 0.16 | 0.58 |

**Supplementary table 5. Parameter estimates of the glutamate secretion per cell**. Associated 95% confidence intervals were computed.

| Cell lines | $k$ | lower CI | upper CI |
|---|---|---|---|
| BJ5ta-WT | 6.94e-06 | 6.09e-06 | 7.78e-06 |
| BJ5ta-MUTKO2 | 8.72e-06 | 7.79e-06 | 9.66e-06 |
| BJ5ta-MUTKO3 | 1.02e-05 | 9.35e-06 | 1.10e-05 |
| BJ5ta-MUTKO7 | 1.09e-05 | 9.71e-06 | 1.20e-05 |

**Supplementary table 6. Difference in estimates of the glutamate secretion per cell**. P-value was adjusted using Dunnett method.

| Cell lines compared | estimate | P-value |
|---|---|---|
| (BJ5ta-MUTKO2) -(BJ5ta-WT) | 6.94e-06 | 0.0214 |
| (BJ5ta-MUTKO3) -(BJ5ta-WT) | 3.26e-06 | $<$0.0001 |
| (BJ5ta-MUTKO7) -(BJ5ta-WT) | 3.93e-06 | $<$0.0001 |

**Supplementary table 7. Parameter estimates of the glutamine consumption per cell**. Associated 95% confidence intervals (CI) were computed.

| Cell lines | $k$ | lower CI | upper CI |
|---|---|---|---|
| BJ5ta-WT | 1.23e-08 | 1.13e-08 | 1.33e-08 |
| BJ5ta-MUTKO2 | 1.64e-08 | 1.52e-08 | 1.76e-08 |
| BJ5ta-MUTKO3 | 2.17e-08 | 2.07e-08 | 2.27e-08 |
| BJ5ta-MUTKO7 | 1.37e-08 | 1.15e-08 | 1.60e-08 |

**Supplementary table 8. Difference in estimates of the glutamine consumption per cell**. P-value was adjusted using Dunnett method. Only the points at 0 and 24h were considered.

| Cell lines compared | estimate | P-value |
|---|---|---|
| (BJ5ta-MUTKO2) -(BJ5ta-WT) | 4.12e-09 | 0.0003 |
| (BJ5ta-MUTKO3) -(BJ5ta-WT) | 9.38e-09 | $<$0.0001 |
| (BJ5ta-MUTKO7) -(BJ5ta-WT) | 1.44e-09 | 0.46 |

**Supplementary table 9. Parameter estimates of the glutamine kill curve**. A 3-parameter logistic curve of the form was fitted: $\frac{b_1}{1+(\frac{t}{b_3})^{b_2}}$. Associated 95% confidence intervals were computed.

| Cell lines | $b_2$ | lower CI | upper CI | $b_3$ | lower CI | upper CI |
|---|---|---|---|---|---|---|
| BJ5ta-WT | 2.60 | 0.43 | 4.77 | 208.67 | 168.65 | 248.68 |
| BJ5ta-MUTKO2 | 2.97 | 1.63 | 4.30 | 138.44 | 113.66 | 163.22 |
| BJ5ta-MUTKO3 | 3.39 | 2.26 | 4.53 | 151.71 | 135.87 | 167.56 |
| BJ5ta-MUTKO7 | 3.94 | 2.11 | 5.76 | 183.41 | 166.62 | 200.21 |

**Supplementary table 10. List of genes with mutations of the BJ5ta cell lines.** M2, M3 and M7 stand for MUTKO-2, MUTKO-3 and MUTKO-7, respectively. Chr stands for chromosome.

| M2 | M3 | M7 | Chr | Position | Ref | Alt | Zygosity | Genes | Description | Variant description | varType |
|---|---|---|---|---|---|---|---|---|---|---|---|
| 1 | | | chr1 | 58054958 | A | G | HETEROZYGOUS:0/1 | RPS26P15 | ribosomal protein S26 pseudogene 15 | Downstream gene variant | SNV |
| 1 | | | chr1 | 152121000 | C | T | HETEROZYGOUS:0/ | AL589986.1 | | Upstream gene variant | SNV |
| | | 1 | chr1 | 152123140 | GA | G | HETEROZYGOUS:0/ | AL589986.1 | | Intron variant and non coding transcript variant | InDel |
| | 1 | | chr1 | 239204005 | AT | A | HETEROZYGOUS:0/ | AC093426.2 | | Upstream gene variant | InDel |
| 1 | | | chr11 | 76755067 | T | C | HETEROZYGOUS:0/ | AP003119.1 | | Downstream gene variant | SNV |
| | | 1 | chr12 | 30316880 | C | A | HETEROZYGOUS:0/ | AC078776.1 | | Upstream gene variant | SNV |
| | | 1 | chr14 | 57836695 | G | T | HETEROZYGOUS:0/ | RN7SKP99 | RN7SK pseudogene 99 | Downstream gene variant | SNV |
| | 1 | | chr14 | 66258045 | G | T | HETEROZYGOUS:0/ | AL157997.1 | | Intron variant and non coding transcript variant | SNV |
| 1 | | | chr16 | 35085208 | G | T | HETEROZYGOUS:0/ | NAMPTP3 | nicotinamide phosphoribosyltransferase pseudogene 3 | Upstream gene variant | SNV |
| 1 | | | chr16 | 73939131 | A | G | HETEROZYGOUS:0/ | RPSAP56 | ribosomal protein SA pseudogene 56 | Downstream gene variant | SNV |
| 1 | | | chr17 | 17035747 | C | A | HETEROZYGOUS:0/ | AC104024.4 | | Intron variant and non coding transcript variant | SNV |
| | 1 | | chr17 | 27281632 | G | A | HETEROZYGOUS:0/ | RPS16P8 | ribosomal protein S16 pseudogene 8 | Downstream gene variant | SNV |
| | 1 | | chr20 | 25730070 | A | G | HETEROZYGOUS:0/ | VN1R108P | vomeronasal 1 receptor 108 pseudogene | Upstream gene variant | SNV |
| 1 | | | chr22 | 22765261 | G | A | HETEROZYGOUS:0/ | IGLV3-13 | immunoglobulin lambda variable 3-13 (pseudogene) | Downstream gene variant | SNV |
| | 1 | | chr3 | 151203114 | C | T | HETEROZYGOUS:0/ | GPR171 | G protein-coupled receptor 171 | 5 prime UTR variant | SNV |
| | | 1 | chr6 | 26246364 | G | T | HETEROZYGOUS:0/ | H3C7 | H3 clustered histone 7 | Downstream gene variant | SNV |
| 1 | 1 | 1 | chr6 | 104025843 | T | C | HETEROZYGOUS:0/ | NPM1P10 | nucleophosmin 1 pseudogene 10 | Non coding transcript exon variant | SNV |
| | | 1 | chr7 | 128864116 | G | T | HETEROZYGOUS:0/ | ATP6V1FNB | ATP6V1F (ATPase H+ transporting V1 subunit F) neighbor | Upstream gene variant | SNV |
| | | 1 | chrX | 13760933 | G | C | HOMOZYGOUS:1/1 | OFD1 | OFD1 centriole and centriolar satellite protein | Intron variant | SNV |
| 1 | | | chrX | 17765540 | T | A | HOMOZYGOUS:1/1 | Z93242.1 | | Downstream gene variant | SNV |
| 1 | 1 | 1 | chrX | 21132109 | TA | T | HOMOZYGOUS:1/1 | BX088723.1 | | Intron variant and non coding transcript variant | InDel |
| | | 1 | chrX | 21151553 | A | C | HOMOZYGOUS:1/1 | BX088723.1 | | Intron variant and non coding transcript variant | SNV |
| 1 | 1 | 1 | chrX | 21166492 | G | T | HOMOZYGOUS:1/1 | BX088723.1 | | Downstream gene variant | SNV |
| | 1 | | chrX | 52765160 | G | A | HOMOZYGOUS:1/1 | SSX2B | SSX family member 2B | Downstream gene variant | SNV |
| | | 1 | chrX | 87430159 | A | T | HOMOZYGOUS:1/1 | Z96811.1 | | Upstream gene variant | SNV |
| | | 1 | chrX | 103348251 | C | A | HOMOZYGOUS:1/1 | Z92846.1 | | Upstream gene variant | SNV |
| 1 | | | chrX | 120867507 | A | C | HOMOZYGOUS:1/1 | PA2G4P1 | proliferation-associated 2G4 pseudogene 1 | Downstream gene variant | SNV |
| | 1 | | chrX | 122245583 | G | C | HOMOZYGOUS:1/1 | AL357562.1 | | Downstream gene variant | SNV |
| 1 | 1 | 1 | chrX | 139443874 | GCCA( | G | HOMOZYGOUS:1/1 | SRD5A1P1 | steroid 5 alpha-reductase 1 pseudogene 1 | Upstream gene variant | InDel |
| 1 | | | chrX | 149605418 | TCTAT | T | HOMOZYGOUS:1/1 | TMEM185A | transmembrane protein 185A | Intron variant | InDel |

### Supplementary table 11. List of reactions predicted to have a different flux according to the sensitivity method

| Reaction ID | Reaction name | Sensitivity |
|---|---|---|
| MMTSADm | malonate-semialdehyde dehydrogenase (acetylating), mitochondrial | -0.53 |
| MMMm | methylmalonyl-CoA mutase | -1.00 |
| MMEm | methylmalonyl-CoA epimerase/racemase | 0.47 |
| MMCDm | Methylmalonyl-CoA decarboxylase, mitochondrial | 0.22 |
| PPCOACm | Propionyl-CoA carboxylase, mitochondrial | -0.21 |
| AKGDm | 2-oxoglutarate dehydrogenase | 0.28 |
| r0384 | 2-oxoglutarate:[dihydrolipoyllysine-residue succinyltransferase]-lipoyllysine 2-oxidoreductase (decarboxylating, acceptor-succinylating) | 0.10 |
| r0620 | 2-oxoglutarate dehydrogenase E1 component Citrate cycle (TCA cycle) | 0.05 |
| r0163 | 2-oxoglutarate dehydrogenase E1 component Citrate cycle (TCA cycle) | 0.05 |
| r0556 | succinyl-CoA:enzyme N6-(dihydrolipoyl)lysine S-succinyltransferase Citrate cycle (TCA cycle) | -0.15 |
| EX_3aib(e) | L-3-Amino-isobutanoate exchange | 0.12 |
| 3AIBTm | L-3-aminoisobutyrate transaminase, mitochondrial | 0.12 |
| 3AIBt | 3-amino-isobutyrate transport | 0.12 |
| 3AIBtm | 3-amino-isobutyrate transport, mitochondrial | 0.12 |
| MMSAD1m | methylmalonate-semialdehyde dehydrogenase | 0.16 |
| 3AIB_Dtm | transport of 3aib_D into mitochondria | -0.17 |
| D3AIBTm | D-3-Amino-isobutanoate:pyruvate aminotransferase, mitochondrial | -0.08 |
| RE3326M | RE3326 | 0.04 |
| r0385 | 3-methyl-2-oxobutanoate:[dihydrolipoyllysine-residue (2-methylpropanoyl)transferase] lipoyllysine 2-oxidoreductase (decarboxylating, acceptor-2-methylpropanoylating) | -0.04 |
| SUCOASm | Succinate–CoA ligase (ADP-forming) | 0.16 |
| ECOAH12m | 3-hydroxyacyl-CoA dehydratase (3-hydroxyisobutyryl-CoA) (mitochondria) | -0.05 |
| 3HBCOAHLm | 3-hydroxyisobutyryl-CoA hydrolase, mitochondrial | -0.05 |
| r0669 | (S)-3-Hydroxyisobutyryl-CoA hydro-lyase | -0.05 |
| r0779 | 3-Hydroxy-2-methylpropanoyl-CoA hydrolase | -0.05 |
| HIBDm | 3-hydroxyisobutyrate dehydrogenase, mitochondrial | -0.11 |
| ECOAH9m | 2-Methylprop-2-enoyl-CoA (2-Methylbut-2-enoyl-CoA), mitochondrial | -0.07 |
| ACACT10m | acetyl-CoA C-acetyltransferase, mitochondrial | -0.07 |
| HACD9m | 3-hydroxyacyl-CoA dehydrogenase (2-Methylacetoacetyl-CoA), mitochondrial | -0.07 |
| 3MOBt2im | 3-methyl-2-oxobutanoate mitochondrial transport via proton symport | -0.06 |
| r0483 | (R)-3-Amino-2-methylpropanoate:2-oxoglutarate aminotransferase | -0.10 |
| ACOAD9m | acyl-CoA dehydrogenase (isobutyryl-CoA), mitochondrial | -0.05 |
| OCOAT1m | 3-oxoacid CoA-transferase | -0.11 |
| r0560 | 2-Methylpropanoyl-CoA:oxygen 2,3-oxidoreductase | -0.06 |
| PPAtm | Propionate transport, diffusion | -0.08 |
| D_3AIBt | D-3-amino-isobutyrate transport | 0.11 |
| EX_3aib_D(e) | D-3-Amino-isobutanoate exchange | 0.11 |
| r0571 | (S)-Methylmalonyl-CoA hydrolase Propanoate metabolism | 0.03 |
| r0643 | (S)-Methylmalonate semialdehyde:NAD+ oxidoreductase | -0.03 |
| SUCOAS1m | Succinate–CoA ligase (GDP-forming) | 0.13 |
| r1154 | EC:1.2.7.2 | -0.13 |
| OIVD2m | 2-oxoisovalerate dehydrogenase (acylating; 3-methyl-2-oxobutanoate), mitochondrial | -0.07 |
| ABTArm | 4-aminobutyrate transaminase, reversible (mitochondrial) | 0.09 |
| 4ABUTtm | 4-aminobutanoate mitochondrial transport via diffusion | 0.09 |
| r0178 | Succinate-semialdehyde:NAD+ oxidoreductase | 0.05 |
| ALASm | 5-aminolevulinate synthase | -0.12 |

| Reaction ID | Reaction name | Sensitivity |
|---|---|---|
| r0196 | Succinyl-CoA:glycine C-succinyl-transferase(decarboxylating) | -0.06 |
| 5AOPtm | 5-Aminolevulinate mitochondrial transport | 0.18 |
| r0517 | Succinyl-CoA:glycine C-succinyl-transferase(decarboxylating) | -0.06 |
| CSNAT2m | carnitine O-aceyltransferase, mitochondrial | -0.06 |
| EX_ppa(e) | Propionate exchange | 0.14 |
| PRPNCOAHYDm | Propenoyl-CoA hydrolase (m) | 0.07 |
| r0365 | 3-hydroxypropanoate:NAD+ oxidoreductase beta-Alanine metabolism | 0.07 |
| r0596 | 3-hydroxyisobutyryl-CoA hydrolase beta-Alanine metabolism | 0.07 |
| r1400 | Active transport | -0.08 |
| r0670 | (S)-3-Methyl-2-oxopentanoate:[dihydrolipoyllysine-residue (2-methylpropanoyl)transferase] lipoyllysine 2-oxidoreductase (decarboxylating, acceptor-2-methylpropanoylating) | -0.02 |
| r0604 | (S)-2-methylbutanoyl-CoA:enzyme N6-(dihydrolipoyl)lysine S-(2-methylbutanoyl)transferase | -0.02 |
| r1291 | Postulated transport reaction | -0.08 |
| r1454 | TCDB:2.A.1.13.5 TCDB:2.A.1.13.1 | 0.07 |
| PCRNtm | transport into the mitochondria from cytosol (carnitine) | -0.03 |
| VALTAm | valine transaminase, mitochondiral | -0.05 |
| VALt5m | Valine reversible mitochondrial transport | -0.05 |
| PPCOAOm | Propanoyl-CoA:FAD 2,3-oxidoreductase, mitochondrial | 0.03 |
| r1155 | TCDB:2.A.1.13.5 TCDB:2.A.1.13.1 | -0.06 |
| EX_succ(e) | Succinate exchange | -0.10 |
| SUCCt4_2 | succinate transport via sodium symport | 0.10 |
| r0683 | Propanoyl-CoA:(acceptor) 2,3-oxidoreductase | 0.03 |
| r2437 | Mitochondrial Carrier (MC) TCDB:2.A.29.8.3 | -0.03 |
| SUCD1m | succinate dehydrogenase | -0.15 |
| ME2m | malic enzyme (NADP), mitochondrial | -0.07 |
| r1659 | Amino Acid-Polyamine-Organocation (APC) TCDB:2.A.3.8.1 | 0.00 |
| r0221 | Propinol adenylate:CoA ligase (AMP-forming) | -0.02 |
| ACCOALm | acetate-CoA ligase (AMP-forming) | -0.04 |
| r0319 | Propanoate:CoA ligase (AMP-forming) | -0.02 |
| r0179 | Succinate-semialdehyde:NADP+ oxidoreductase | 0.04 |
| FAOXC6DCC4DCx | fatty acid beta oxidation(C6DC–>C4DC)x | 0.05 |
| r2122 | Major Facilitator(MFS) TCDB:2.A.1.13.1 | -0.01 |
| r0603 | (S)-2-methylbutanoyl-CoA:acceptor 2,3-oxidoreductase | -0.02 |
| FAOXC8060m | R_FAOXC8060m | 0.04 |
| FUMm | fumarase, mitochondrial | -0.08 |
| ME1m | malic enzyme (NAD), mitochondrial | -0.06 |
| PCm | pyruvate carboxylase | 0.07 |
| CSNAT3x | carnitine O-aceyltransferase, peroxisomal | -0.03 |
| SUCCt2m | succinate transport, mitochondrial | 0.04 |
| r2123 | Major Facilitator(MFS) TCDB:2.A.1.13.1 | -0.01 |
| FAOXC6040m | R_FAOXC6040m | 0.03 |
| DHPM2 | dihydropyrimidinase (dihydrothymine) | -0.06 |
| BUP2 | beta-ureidopropionase (D-3-amino-isobutanoate forming) | -0.06 |
| r2121 | Major Facilitator(MFS) TCDB:2.A.1.13.1 | -0.01 |
| r2103 | Major Facilitator(MFS) TCDB:2.A.1.13.1 | 0.02 |
| r2382 | Mitochondrial Carrier (MC) TCDB:2.A.29.7.2 | -0.01 |
| r2096 | Major Facilitator(MFS) TCDB:2.A.1.13.1 | 0.01 |
| adpact | adpact | 0.02 |
| EX_adpac | EX_adpac | -0.02 |
| r0321 | Acetoacetate:CoA ligase (AMP-forming) | 0.04 |
| C4DCCACT | transport of succinyl carnitine into cytosol | 0.02 |
| SUCCCROT | production of succinyl carnitine | 0.02 |
| FAOXC4C4DCc | succ–>C4DCc | -0.02 |
| SUCCACT | activation of succinate | -0.02 |

| Reaction ID | Reaction name | Sensitivity |
|---|---|---|
| FUMtm | fumarate transport, mitochondrial | 0.04 |
| r2124 | Major Facilitator(MFS) TCDB:2.A.1.13.1 | -0.01 |
| SUCCTD | transport of succinate by diffusion | 0.04 |
| EX_2hb(e) | 2-Hydroxybutyrate exchange | 0.06 |
| 2HBO | 2-Hydroxybutyrate:NAD+ oxidoreductase | -0.06 |
| FACOAL150 | fatty-acid–CoA ligase | -0.01 |
| PTDCAt | fatty acid transport via diffusion | -0.01 |
| EX_ptdca(e) | pentadecanoate exchange | 0.01 |
| PTDCACRNCPT1 | carnitine fatty-acyl transferase | -0.01 |
| PTDCACRNCPT2 | pentadecanoate transport into the mitochondria | -0.01 |
| PTDCACRNt | pentadecanoate transport into the mitochondria | -0.01 |
| GLYtm | glycine passive transport to mitochondria | -0.06 |
| 3MOBte | Transport of 3-methyl-2-oxobutanoate | -0.05 |
| EX_3mob(e) | Exchange of 3-methyl-2-oxobutanoate | 0.05 |
| SUCCOAPET | thioesterification of succinyl coa for release into cytosol | 0.03 |
| FPGS2m | folylpolyglutamate synthetase, mitochondrial | -0.01 |
| r0557 | Glutaryl-CoA:dihydrolipoamide S-succinyltransferase | -0.02 |
| r0451 | 2-Oxoadipate:lipoamde 2-oxidoreductase(decarboxylating and acceptor-succinylating) | -0.02 |
| r0555 | acetyl-CoA:enzyme N6-(dihydrolipoyl)lysine S-acetyltransferase | -0.04 |
| r0383 | pyruvate:[dihydrolipoyllysine-residue acetyltransferase]-lipoyllysine 2-oxidoreductase (decarboxylating, acceptor-acetylating) | -0.04 |
| FAOXCPRIST3x | R_FAOXCPRIST3x | -0.01 |
| FAOXC15BRC13BRx | fatty acid beta oxidation(C15br–>C13br)x | -0.01 |
| FAOXCPRIST2x | R_FAOXCPRIST2x | -0.01 |
| FAOXC13BRC11BRx | fatty acid beta oxidation(C13br–>C11br)x | -0.01 |
| FAOXC11BRC9BRx | fatty acid beta oxidation(C11br–>C9br)x | -0.01 |
| FAOXCPRIST1x | R_FAOXCPRIST1x | -0.01 |
| DMHPTCRNt | 2,6 dimethylheptanoyl crn transport | 0.02 |
| DMHPTCRNCPT2 | 2,6 dimethylheptanoly CoA carnitine transferase | 0.02 |
| DMNONCRNCPT2 | DMNONCRNCPT2 | -0.02 |
| EX_dmhptcrn(e) | 2,6 dimethylheptanoyl carnitine exchange | -0.02 |
| DMHPTCRNte | 2,6 dimethylheptanoyl carnitine transport | -0.02 |
| FAOXC9BRC7BRm | fatty acid beta oxidation(C9br–>C7br)m | -0.02 |
| r0295 | glycine synthase | -0.03 |
| GCCbim | glycine-cleavage complex (lipoylprotein) irreversible, mitochondrial | 0.01 |
| GCCam | glycine-cleavage complex (lipoylprotein), mitochondrial | 0.01 |
| GCCcm | glycine-cleavage complex (lipoylprotein), mitochondrial | 0.01 |
| C30CPT1 | production of propionylcarnitine | -0.03 |
| r0975 | Facilitated diffusion | -0.02 |
| r0331 | 5,6-Dihydrothymine:NAD+ oxidoreductase | -0.02 |
| r0921 | Postulated transport reaction | -0.02 |
| ACOAD10m | acyl-CoA dehydrogenase (2-methylbutanoyl-CoA), mitochondrial | -0.02 |
| r2111 | Major Facilitator(MFS) TCDB:2.A.1.13.1 | 0.01 |
| HMGCOASim | Hydroxymethylglutaryl CoA synthase (ir) | -0.04 |
| GCC2cm | glycine-cleavage complex (lipoamide), mitochondrial | 0.02 |
| EX_prist | EX_prist | 0.03 |
| PRISTt | PRISTt | -0.03 |
| CRNCAR3tp | carnitine-propcarnitine carrier, peroxisomal | -0.02 |
| DMNONCRNt | 4,8 dimethylnonanoyl carnitine transport (mitochondria) | -0.01 |
| r0636 | Octanoyl-CoA:L-carnitine O-octanoyltransferase EC:2.3.1.137 | -0.02 |
| r2434 | Mitochondrial Carrier (MC) TCDB:2.A.29.8.3 | -0.02 |
| THYMt | thymine reversible transport via facilated diffusion | -0.03 |

| Reaction ID | Reaction name | Sensitivity |
|---|---|---|
| EX_thym(e) | Thymine exchange | 0.03 |
| FPGSm | folylpolyglutamate synthetase, mitochondrial | -0.02 |
| r0834 | Mitochondrial Carrier (MC) TCDB:2.A.29.2.2 | 0.02 |
| CRNtx | carnitine transport peroxisome to mitochondria | -0.01 |
| LDH_Lm | L-lactate dehydrogenase | 0.03 |
| FE2tm | iron (II) transport | -0.02 |
| PPPG9tm | protoporphyrinogen IX mitochondrial transport | -0.02 |
| PHEMEtm | Heme transport to cytosol | -0.02 |
| FCLTm | Ferrochelatase, mitochondrial | -0.02 |
| PPPGOm | protoporphyrinogen oxidase, mitochondrial | -0.01 |
| CPPPGO | coproporphyrinogen oxidase (O2 required) | -0.02 |
| HMBS | hydroxymethylbilane synthase | -0.02 |
| PPBNGS | porphobilinogen synthase | -0.07 |
| UPP3S | uroporphyrinogen-III synthase | -0.02 |
| UPPDC1 | uroporphyrinogen decarboxylase (uroporphyrinogen III) | -0.02 |
| 2MCITt | 2-methylcitrate transport via diffusion | 0.05 |
| MCITS | 2-methylcitrate synthase | 0.05 |
| EX_2mcit(e) | 2-Methylcitrate exchange | 0.05 |
| r2133 | Major Facilitator(MFS) TCDB:2.A.1.13.1 | 0.01 |
| r2378 | Mitochondrial Carrier (MC) TCDB:2.A.29.7.2 | 0.01 |
| r0915 | Mitochondrial Carrier (MC) TCDB:2.A.29.7.2 | -0.02 |
| r2116 | Major Facilitator(MFS) TCDB:2.A.1.13.1 | 0.01 |
| r2390 | Mitochondrial Carrier (MC) TCDB:2.A.29.7.2 | -0.01 |
| FPGS6m | folylpolyglutamate synthetase (DHF), mitochondrial | -0.01 |
| 7DHFtm | 7-glutamyl-DHF transport,m mitochondrial | -0.01 |
| EX_4abut(e) | 4-Aminobutanoate exchange | -0.04 |
| THRGLNexR | L-threonine/glycine reversible exchange | 0.00 |
| 7THFtm | 7-glutamyl-THF transport,m mitochondrial | -0.01 |
| FPGS3m | folylpolyglutamate synthetase, mitochondrial | -0.01 |
| ILETAm | isoleucine transaminase, mitochondrial | -0.02 |
| ILEt5m | Isoleucine mitochondrial transport | -0.02 |
| ARGDCm | arginine decarboxylase (m) | -0.02 |
| r0907 | Active transport | -0.02 |
| AGMTm | agmatinase (m) | -0.02 |
| NDPK1m | nucleoside-diphosphate kinase (ATP:GDP), mitochondrial | 0.02 |
| ITCOALm | Itaconate–CoA ligase (ADP-forming), mitochondrial | 0.01 |
| MECOAS1m | mesaconate–CoA ligase (GDP-forming) | -0.01 |
| MECOALm | mesaconate–CoA ligase (ADP-forming), mitochondrial | 0.01 |
| ITCOAL1m | Itaconate–CoA ligase (GDP-forming), mitochondrial | -0.01 |
| 3MOPt2im | 3-Methyl-2-oxopentanoate mitochondrial transport via proton symport | -0.02 |
| CYSTGL | cystathionine g-lyase | -0.05 |
| r2420 | Mitochondrial Carrier (MC) TCDB:2.A.29.2.7 | 0.02 |
| EX_val_L(e) | L-Valine exchange | 0.07 |
| r2520 | Mitochondrial Carrier (MC) TCDB:2.A.29.2.1 | -0.03 |
| L_LACtm | L-lactate transport, mitochondrial | 0.02 |
| FAOXC8DCC6DCx | fatty acid beta oxidation(C8DC–>C6DC)x | 0.03 |
| r1109 | Citrate oxaloacetate-lyase ((pro-3S)-CH2COO- –>acetate) | -0.04 |
| 6THFtm | 6-glutamyl-THF transport,m mitochondrial | 0.00 |
| FAOXC5030m | R_FAOXC5030m | -0.01 |
| FAOXC7050m | R_FAOXC7050m | -0.01 |
| FAOXC7C5m | fatty acid beta oxidation(C7–>C5)m | -0.01 |
| FAOXC15C13m | fatty acid beta oxidation(C15–>C13)m | -0.01 |
| FAOXC9070m | R_FAOXC9070m | -0.01 |
| FAOXC11C9m | fatty acid beta oxidation(C11–>C9)m | -0.01 |
| FAOXC11090m | R_FAOXC11090m | -0.01 |
| FAOXC13C11m | fatty acid beta oxidation(C13–>C11)m | -0.01 |

| Reaction ID | Reaction name | Sensitivity |
|---|---|---|
| FAOXC5C3x | fatty acid beta oxidation(C5–>C3)m | -0.01 |
| FAOXC9C7m | fatty acid beta oxidation(C9–>C7)m | -0.01 |
| FAOXC130110m | R_FAOXC130110m | -0.01 |
| FAOXC150130m | R_FAOXC150130m | -0.01 |
| r0822 | Mitochondrial Carrier (MC) TCDB:2.A.29.2.2 | 0.01 |
| 10FTHF7GLUtm | 7-glutamyl-10FTHF transport, mitochondrial | -0.01 |
| FPGS9m | folylpolyglutamate synthetase (10fthf), mitochondrial | -0.01 |
| r1649 | Amino Acid-Polyamine-Organocation (APC) TCDB:2.A.3.8.1 | 0.00 |
| r2090 | Major Facilitator(MFS) TCDB:2.A.1.13.1 | 0.01 |
| COAtm | CoA transporter | 0.05 |
| FPGS8m | folylpolyglutamate synthetase (10fthf), mitochondrial | -0.01 |
| EX_c51crn_ | exchange reaction for tiglyl carnitine | 0.02 |
| TIGCRNe | transport of tiglyl carnitine into the extra cellular fluid | 0.02 |
| C51CPT1 | production of tiglylcarnitine | 0.02 |
| 2MB2COAc | transport of 2-methylcrotonoyl-CoA into cytosol | 0.02 |
| r0714 | (S)-3-Hydroxyhexadecanoyl-CoA:NAD+ oxidoreductase | 0.00 |
| r0653 | myristoyl-CoA:acetylCoA C-myristoyltransferase | 0.00 |
| TMDPP | thymidine phosphorylase | -0.03 |
| r2091 | Major Facilitator(MFS) TCDB:2.A.1.13.1 | 0.01 |
| OIVD3m | 2-oxoisovalerate dehydrogenase (acylating; 3-methyl-2-oxopentanoate), mitochondrial | -0.02 |
| r1450 | EC:1.3.3.6 | 0.01 |
| DM_4abut(n) | Demand for 4-Aminobutanoate(n) | -0.03 |
| 4ABUTtcn | transport of GABA | -0.03 |
| L_LACtcm | L-lactate transport via diffusion (cytosol to mitochondria) | 0.01 |
| SARDHm | Sarcosine dehydrogenase (m) | -0.03 |
| r2375 | Mitochondrial Carrier (MC) TCDB:2.A.29.7.2 | 0.02 |
| SARCStm | Sarcosine transport (mitochondrial) | -0.03 |
| r2082 | Major Facilitator(MFS) TCDB:2.A.1.13.1 | -0.01 |
| MMSAD3m | methylmalonate-semialdehyde dehydrogenase (malonic semialdehyde), mitochondrial | 0.04 |
| FAOXC141_5Zm | R_FAOXC141_5Zm | 0.01 |
| FAOXC121_3Zm | R_FAOXC121_3Zm | 0.01 |
| FAOXC181_9Zm | R_FAOXC181_9Zm | 0.01 |
| FAOXC161_7Zm | R_FAOXC161_7Zm | 0.01 |
| ADPACTD | transport into cytosol (diffusion) | -0.02 |
| ADPCOAPTE | thioesterification of adipoyl coA for release into cytosol | -0.02 |
| CBPSam | carbamoyl-phosphate synthase (ammonia) (mitochondria | 0.04 |
| r1581 | Amino Acid-Polyamine-Organocation (APC) TCDB:2.A.3.8.1 | 0.00 |
| PROtm | L-proline transport, mitochondrial | 0.04 |
| APAT2rm | 3-Aminopropanoate:2-oxoglutarate aminotransferase (m) | -0.03 |
| BALAtmr | Beta-alanine reversible mitochondrial transport (diffusion) | -0.03 |
| ADK3m | adentylate kinase (GTP) | -0.03 |
| subeact | subeact | 0.01 |
| EX_subeac | EX_subeac | -0.01 |
| FAOXC101_3Em | R_FAOXC101_3Em | 0.01 |
| r0731 | (S)-Hydroxyoctanoyl-CoA hydro-lyase | 0.01 |
| r0730 | (S)-Hydroxyoctanoyl-CoA:NAD+ oxidoreductase | 0.01 |
| r0732 | Hexanoyl-CoA:acetyl-CoA C-acyltransferase | 0.01 |
| FAOXC61C4m | fatty acid beta oxidation(C6:1–>C4:0)m | 0.02 |
| r0287 | Acetyl-CoA:acetyl-CoA C-acetyltransferase | 0.01 |
| r0733 | (S)-Hydroxyhexanoyl-CoA:NAD+ oxidoreductase | 0.01 |
| r0734 | (S)-Hydroxyhexanoyl-CoA hydro-lyase | 0.01 |
| HDECAACBP | transport of 3-hydroxyhexadecanoylcoa from mitochondria into the cytosol | -0.01 |
| C16OHc | production of 3-hydroxyhexadecanoylcarnitinec | -0.01 |
| HEXDCRNe | transport of 3-hydroxyhexadecanoyl carnitine into extracellular space | -0.01 |
| EX_3hexdcrn_ | exchange reaction for 3-hydroxyhexadecanoyl carnitine | -0.01 |

| Reaction ID | Reaction name | Sensitivity |
|---|---|---|
| r0639 | Lauroyl-CoA:acetyl-CoA C-acyltransferase | 0.01 |
| r0718 | (S)-3-Hydroxytetradecanoyl-CoA:NAD+ oxidoreductase | 0.01 |
| r1631 | Amino Acid-Polyamine-Organocation (APC) TCDB:2.A.3.8.1 | 0.00 |
| r2386 | Mitochondrial Carrier (MC) TCDB:2.A.29.7.2 | -0.01 |
| r0386 | 4-methyl-2-oxopentanoate:[dihydrolipoyllysine-residue (2-methylpropanoyl)transferase] lipoyllysine 2-oxidoreductase (decarboxylating, acceptor-2-methylpropanoylating) | -0.01 |
| r0656 | 3-methylbutanoyl-CoA:enzyme N6-(dihydrolipoyl)lysine S-(3-methylbutanoyl)transferase | -0.01 |
| r0801 | Mitochondrial Carrier (MC) TCDB:2.A.29.21.1 | 0.03 |
| C60CRNt | C160 transport into the mitochondria | -0.01 |
| C60CPT2 | carnitine O-hexanoyl transferase | -0.01 |
| H2CO3Dm | carboxylic acid dissociation | -0.03 |
| P5CDm | 1-pyrroline-5-carboxylate dehydrogenase, mitochondrial | 0.03 |
| r2109 | Major Facilitator(MFS) TCDB:2.A.1.13.1 | 0.01 |
| FAOXC1811601m | Beta oxidation fatty acid | 0.02 |
| CSNAT2x | carnitine dimethyl nonanoyl transferase, revsible, peroxisomal | -0.01 |
| CO2tm | CO2 transport (diffusion), mitochondrial | -0.07 |
| FAOXC140120m | R_FAOXC140120m | 0.01 |
| r0541 | Glutaryl-CoA:(acceptor) 2,3-oxidoreductase (decarboxylating) | -0.02 |
| r0450 | L-2-Aminoadipate:2-oxoglutarate aminotransferase | -0.02 |
| FPGS | folylpolyglutamate synthetase | 0.02 |
| r0074 | L-Glutamate 5-semialdehyde:NAD+ oxidoreductase | 0.03 |
| ACOAD1fm | acyl-CoA dehydrogenase (butanoyl-CoA), mitochondrial | 0.02 |
| r1658 | Amino Acid-Polyamine-Organocation (APC) TCDB:2.A.3.8.1 | 0.00 |
| MALtm | malate transport, mitochondrial | 0.02 |
| C4CRNCPT2 | transport of butyryl carnitine in the mitochondrial matrix for final hydrolysis | -0.01 |
| EX_leuktrF4(e) | leukotriene F4 exchange | -0.02 |
| AGTim | alanine-glyoxylate transaminase (irreversible), mitochondrial | -0.03 |
| FAOXC141C121m | fatty acid beta oxidation(C14:1–>C12:1)m | 0.01 |
| FAOXC121C101m | fatty acid beta oxidation(C12:1–>C10:1)m | 0.01 |
| THRD_L | L-threonine deaminase | -0.06 |
| ELAIDCPT2 | carnitine transferase | 0.02 |
| PCRNtc | transport into the cytosol from peroxisome (carnitine) | -0.01 |
| GLUt2m | L-glutamate reversible transport via proton symport, mitochondrial | 0.05 |
| FAOXC160140m | R_FAOXC160140m | 0.01 |
| FPGS2 | folylpolyglutamate synthetase | 0.01 |
| C181CPT2 | transport of Octadecenoyl-CoA into mitochondrial matrix | -0.01 |
| r2419 | Mitochondrial Carrier (MC) TCDB:2.A.29.2.7 | 0.03 |
| EX_thr_L(e) | L-Threonine exchange | 0.06 |
| r0081 | L-Alanine:2-oxoglutarate aminotransferase | -0.02 |
| PDHm | pyruvate dehydrogenase | -0.06 |
| UREAtm | Urea transport via diffusion | 0.02 |
| VALLAT1tc | transport of L-Valine by LAT1 in association with 4F2hc, across the apical surface of the memebranes. | 0.00 |
| EX_cyst_L | EX_cyst_L | 0.03 |
| cyst_Lt | cyst_Lt | -0.03 |
| r1654 | Amino Acid-Polyamine-Organocation (APC) TCDB:2.A.3.8.1 | 0.00 |
| FAOXC18C18OHm | fatty acid beta oxidation(C18–>C18OH)m | -0.01 |
| C18OHc | production of 3-hydroxyoctadecanoylcarnitine | -0.01 |
| HOCDACBP | transport of (S)-3-Hydroxyoctadecanoyl-CoA from mitochondria into the cytosol | -0.01 |
| HOCTDECCRNe | transport of 3-hydroxyoctadecanoyl carnitine into extracellular space | -0.01 |
| EX_3octdeccrn_ | exchange reaction for 3-hydroxyoctadecanoyl carnitine | -0.01 |
| FPGS4m | folylpolyglutamate synthetase (DHF), mitochondrial | -0.01 |
| FPGS5m | folylpolyglutamate synthetase (DHF), mitochondrial | -0.01 |

| Reaction ID | Reaction name | Sensitivity |
|---|---|---|
| FAOXC170150m | R_FAOXC170150m | -0.01 |
| HPDCAt | fatty acid transport via diffusion | -0.01 |
| EX_hpdca(e) | heptadecanoate exchange | 0.01 |
| HPDCACRNCPT2 | heptadecanoate transport into the mitochondria | -0.01 |
| HPDCACRNt | heptadecanoate transport into the mitochondria | -0.01 |
| FACOAL170 | fatty-acid–CoA ligase | -0.01 |
| HPDCACRNCPT1 | carnitine fatty-acyl transferase | -0.01 |
| r2086 | Major Facilitator(MFS) TCDB:2.A.1.13.1 | 0.01 |
| C181OHc | production of 3-hydroxyoctadecenoylcarnitinec | -0.01 |
| HOCTDACBP | transport of 3-hydroxyoctadecenoylcoa from mitochondria into the cytosol | -0.01 |
| FAOXC181C181OHm | fatty acid beta oxidation(C18:1–>C18:1OH)m | -0.01 |
| EX_3octdece1crn_ | exchange reaction for 3-hydroxy-octadecenoyl carnitine | -0.01 |
| OCTDECE1CRNe | transport of 3-hydroxy-octadecenoyl carnitine into extracellular space | -0.01 |
| C100CPT2 | carnitine O-hexanoyl transferase | -0.01 |
| C100CRNt | C160 transport into the mitochondria | -0.01 |
| FAOXC120100m | R_FAOXC120100m | 0.02 |
| r0724 | Decanoyl-CoA:acetyl-CoA C-acyltransferase | -0.01 |
| r0722 | (S)-3-Hydroxydodecanoyl-CoA:NAD+ oxidoreductase | 0.01 |
| H2Otm | H2O transport, mitochondrial | 0.04 |
| r1650 | Amino Acid-Polyamine-Organocation (APC) TCDB:2.A.3.8.1 | 0.00 |
| r1660 | Amino Acid-Polyamine-Organocation (APC) TCDB:2.A.3.8.1 | 0.00 |
| GLUTCOADHm | glutaryl-CoA dehydrogenase (mitochondria) | -0.02 |
| EHGLATm | L-erythro-4-Hydroxyglutamate:2-oxoglutarate aminotransferase, mitochondrial | -0.02 |
| FAOXC180 | fatty acid beta oxidation(C18–>C16)m | 0.02 |
| ACCOACm | Acetyl-Coa carboxylase, beta isoform | 0.02 |
| FAOXC141_7Em | R_FAOXC141_7Em | 0.01 |
| FAOXC121_5Em | R_FAOXC121_5Em | 0.01 |
| FAOXC161_9Em | R_FAOXC161_9Em | 0.01 |
| FAOXC181_11Em | R_FAOXC181_11Em | 0.01 |
| r1464 | Active transport | -0.03 |
| r0726 | (S)-Hydroxydecanoyl-CoA:NAD+ oxidoreductase | 0.01 |
| r0634 | Octanoyl-CoA:acetyl-CoA C-acyltransferase | -0.01 |
| r0941 | Free diffusion | -0.04 |
| C14OHc | production of 3-hydroxytetradecanoylcarnitine | -0.01 |
| EX_3tdcrn_ | exchange reaction for 3-hydroxy-tetradecanoyl carnitine | -0.01 |
| HTDCRNe | transport of 3-hydroxy-tetradecanoyl carnitine into extracellular space | -0.01 |
| HTDCACBP | transport of (S)-3-Hydroxytetradecanoyl-CoA from mitochondria into the cytosol | -0.01 |
| r1434 | Transport reaction | 0.02 |
| EX_akg(e) | 2-Oxoglutarate exchange | -0.05 |
| EX_leuktrD4(e) | leukotriene D4 exchange | 0.02 |
| SCP22x | Sterol carrier protein 2 | 0.01 |
| DMNONCOACRNCP | DMNONCOACRNCPT1 | -0.01 |
| r0835 | Mitochondrial Carrier (MC) TCDB:2.A.29.2.2 | -0.01 |
| r1614 | Amino Acid-Polyamine-Organocation (APC) TCDB:2.A.3.8.1 | 0.00 |
| r2089 | Major Facilitator(MFS) TCDB:2.A.1.13.1 | -0.01 |
| DHFtm | dihydrofolate reversible mitochondrial transport | -0.01 |
| 5aopt | 5aopt | 0.05 |
| EX_5aop | EX_5aop | -0.05 |
| PRO1xm | proline oxidase (NAD), mitochondrial | 0.02 |
| EX_HC00955(e) | L-3-Cyanoalanine exchange | 0.02 |
| EX_c8crn_ | exchange reaction for octanoyl carnitine | 0.01 |
| C8CRNe | transport of octanoyl carnitine into the extra cellular space | 0.01 |
| r1446 | EC:1.3.3.6 | 0.02 |

| Reaction ID | Reaction name | Sensitivity |
|---|---|---|
| RE2649M | RE2649 | 0.02 |
| r1653 | Amino Acid-Polyamine-Organocation (APC) TCDB:2.A.3.8.1 | 0.00 |
| LPCOXp | L-pipecolate oxidase, peroxisomal | -0.01 |
| RE1254C | RE1254 | -0.01 |
| THP2Ctp | 2,3,4,5-Tetrahydropyridine-2-carboxylate transport, peroxisomal | 0.01 |
| r0330 | 5,6-Dihydrothymine:NAD+ oxidoreductase | -0.02 |
| r0947 | Mitochondrial Carrier (MC) TCDB:2.A.29.19.1 | -0.01 |
| FAOXC10080m | R_FAOXC10080m | 0.01 |
| ASPGLUm | aspartate-glutamate mitochondrial shuttle | 0.02 |
| ASPTAm | aspartate transaminase | -0.02 |
| GTHRDt | Glutathione transport into mitochondria | 0.03 |
| r1652 | Amino Acid-Polyamine-Organocation (APC) TCDB:2.A.3.8.1 | 0.00 |
| r0830 | Mitochondrial Carrier (MC) TCDB:2.A.29.2.2 | 0.01 |
| CSm | citrate synthase | 0.03 |
| EX_glu_L(e) | L-Glutamate exchange | -0.05 |
| r2087 | Major Facilitator(MFS) TCDB:2.A.1.13.1 | 0.01 |

### Supplementary table 12. *in silico* gene deletion experiment

| Gene Entrez ID | HGNC | Reactions | Gene description | FBA ratio |
|---|---|---|---|---|
| 1738.1 | HGNC:2898 | 2OXOADOXm, AKGDm, GCC2am, GCC2bim, GCC2cm, GCCam, GCCbim, GCCcm, PDHm, r0295, r1154 | dihydrolipoamide dehydrogenase | 0.99 |
| 26275.1 | HGNC:4908 | r0596, r0779, 3HBCOAHLm | 3-hydroxyisobutyryl-CoA hydrolase | 0.95 |
| 32.1 | HGNC:85 | ACCOACm | acetyl-CoA carboxylase beta | 0.99 |
| 1374.1 | HGNC:2328 | r0430, r0431 | carnitine palmitoyltransferase 1A | 0.99 |
| 1376.1 | HGNC:2330 | ADRNCPT2, ARACHCPT2, C160CPT2, C161CPT2, C161CPT22, C180CPT2, C181CPT2, C204CPT2, C226CPT2, DCSPTN1CPT2, DLNLCGCPT2, EICOSTETCPT2, ELAIDCPT2, HPDCACRNCPT2, LNELDCCPT2, LNLCCPT2, LNLNCACPT2, LNLNCGCPT2, PTDCACRNCPT2, RTOTALCRNCPT2, STRDNCCPT2, TMNDNCCPT2, TTDCPT2, VACCCPT2, r0430, r0431, C4CRNCPT2, OCTDECCPT2, C60CPT2, C100CPT2, C120CPT2 | carnitine palmitoyltransferase 2 | 0.99 |
| 1491.1 | HGNC:2501 | CYSTGL, r0193 | cystathionine gamma-lyase | 0.99 |
| 10841.1 | HGNC:3974 | FTCD, GluForTx | formimidoyltransferase cyclodeaminase | 0.99 |
| 2271.1 | HGNC:3700 | FUM, FUMm | fumarate hydratase | 0.99 |
| 4329.1 | HGNC:7179 | MMSAD1m, MMSAD3m | aldehyde dehydrogenase 6 family member A1 | 0.95 |
| 3034.1 | HGNC:4806 | HISD | histidine ammonia-lyase | 0.99 |
| 144193.1 | HGNC:28577 | IZPN | amidohydrolase domain containing 1 | 0.99 |
| 1737.1 | HGNC:2896 | PDHm, r0555 | dihydrolipoamide S-acetyltransferase | 0.99 |
| 6389.1 | HGNC:10680 | SUCD1m | succinate dehydrogenase complex flavoprotein subunit A | 0.99 |
| 6390.1 | HGNC:10681 | SUCD1m | succinate dehydrogenase complex iron sulfur subunit B | 0.99 |
| 6391.1 | HGNC:10682 | SUCD1m | succinate dehydrogenase complex subunit C | 0.99 |
| 6392.1 | HGNC:10683 | SUCD1m | succinate dehydrogenase complex subunit D | 0.99 |
| 131669.1 | HGNC:26444 | URCN | urocanate hydratase 1 | 0.99 |
| 10632.1 | HGNC:14247 | ATPS4m | ATP synthase membrane subunit g | 0.97 |
| 4905.1 | HGNC:8016 | ATPS4m | N-ethylmaleimide sensitive factor, vesicle fusing ATPase | 0.97 |
| 498.1 | HGNC:823 | ATPS4m | ATP synthase F1 subunit alpha | 0.97 |
| 506.1 | HGNC:830 | ATPS4m | ATP synthase F1 subunit beta | 0.97 |
| 509.1 | HGNC:833 | ATPS4m | ATP synthase F1 subunit gamma | 0.97 |
| 513.1 | HGNC:837 | ATPS4m | ATP synthase F1 subunit delta | 0.97 |
| 514.1 | HGNC:838 | ATPS4m | ATP synthase F1 subunit epsilon | 0.97 |
| 515.1 | HGNC:840 | ATPS4m | ATP synthase peripheral stalk-membrane subunit b | 0.97 |
| 10476.1 | HGNC:845 | ATPS4m | ATP synthase peripheral stalk subunit d | 0.97 |
| 521.1 | HGNC:846 | ATPS4m | ATP synthase membrane subunit e | 0.97 |
| 522.1 | HGNC:847 | ATPS4m | ATP synthase peripheral stalk subunit F6 | 0.97 |
| 9551.1 | HGNC:848 | ATPS4m | ATP synthase membrane subunit f | 0.97 |
| 7381.1 | HGNC:12582 | CYOOm2, CYOR_u10m | ubiquinol-cytochrome c reductase binding protein | 0.98 |
| 7384.1 | HGNC:12585 | CYOOm2, CYOR_u10m | ubiquinol-cytochrome c reductase core protein 1 | 0.98 |
| 7385.1 | HGNC:12586 | CYOOm2, CYOR_u10m | ubiquinol-cytochrome c reductase core protein 2 | 0.98 |
| 7386.1 | HGNC:12587 | CYOOm2, CYOR_u10m | ubiquinol-cytochrome c reductase, Rieske iron-sulfur polypeptide 1 | 0.98 |
| 7388.1 | HGNC:12590 | CYOOm2, CYOR_u10m | ubiquinol-cytochrome c reductase hinge protein | 0.98 |
| 1327.1 | HGNC:2265 | CYOOm3 | cytochrome c oxidase subunit 4I1 | 0.99 |
| 9377.1 | HGNC:2267 | CYOOm3 | cytochrome c oxidase subunit 5A | 0.99 |
| 1329.1 | HGNC:2269 | CYOOm3 | cytochrome c oxidase subunit 5B | 0.99 |
| 1337.1 | HGNC:2277 | CYOOm3 | cytochrome c oxidase subunit 6A1 | 0.99 |
| 1339.1 | HGNC:2279 | CYOOm3 | cytochrome c oxidase subunit 6A2 | 0.99 |
| 1340.1 | HGNC:2280 | CYOOm3 | cytochrome c oxidase subunit 6B1 | 0.99 |
| 1345.1 | HGNC:2285 | CYOOm3 | cytochrome c oxidase subunit 6C | 0.99 |
| 1347.1 | HGNC:2288 | CYOOm3 | cytochrome c oxidase subunit 7A2 | 0.99 |
| 1349.1 | HGNC:2291 | CYOOm3 | cytochrome c oxidase subunit 7B | 0.99 |
| 1350.1 | HGNC:2292 | CYOOm3 | cytochrome c oxidase subunit 7C | 0.99 |
| 1351.1 | HGNC:2294 | CYOOm3 | cytochrome c oxidase subunit 8A | 0.99 |
| 170712.1 | HGNC:24381 | CYOOm3 | cytochrome c oxidase subunit 7B2 | 0.99 |
| 341947.1 | HGNC:24382 | CYOOm3 | cytochrome c oxidase subunit 8C | 0.99 |
| 1537.1 | HGNC:2579 | CYOOm2, CYOR_u10m | cytochrome c1 | 0.98 |

| Gene Entrez ID | HGNC | Reactions | Gene description | FBA ratio |
|---|---|---|---|---|
| 27089.1 | HGNC:29594 | CYOOm2, CYOR_u10m | ubiquinol-cytochrome c reductase complex III subunit VII | 0.98 |
| 10975.1 | HGNC:30862 | CYOOm2, CYOR_u10m | ubiquinol-cytochrome c reductase, complex III subunit XI | 0.98 |
| 29796.1 | HGNC:30863 | CYOOm2, CYOR_u10m | ubiquinol-cytochrome c reductase, complex III subunit X | 0.98 |
| 4519.1 | HGNC:7427 | CYOOm2, CYOR_u10m | mitochondrially encoded cytochrome b | 0.98 |
| 51079.1 | HGNC:17194 | NADH2_u10m | NADH:ubiquinone oxidoreductase subunit A13 | 0.99 |
| 126328.1 | HGNC:20371 | NADH2_u10m | NADH:ubiquinone oxidoreductase subunit A11 | 0.99 |
| 55967.1 | HGNC:23987 | NADH2_u10m | NADH:ubiquinone oxidoreductase subunit A12 | 0.99 |
| 7991.1 | HGNC:30242 | NADH2_u10m | tumor suppressor candidate 3 | 0.99 |
| 4535.1 | HGNC:7455 | NADH2_u10m | mitochondrially encoded NADH dehydrogenase 1 | 0.99 |
| 4536.1 | HGNC:7456 | NADH2_u10m | mitochondrially encoded NADH dehydrogenase 2 | 0.99 |
| 4537.1 | HGNC:7458 | NADH2_u10m | mitochondrially encoded NADH dehydrogenase 3 | 0.99 |
| 4538.1 | HGNC:7459 | NADH2_u10m | mitochondrially encoded NADH dehydrogenase 4 | 0.99 |
| 4539.1 | HGNC:7460 | NADH2_u10m | mitochondrially encoded NADH 4L dehydrogenase | 0.99 |
| 4540.1 | HGNC:7461 | NADH2_u10m | mitochondrially encoded NADH dehydrogenase 5 | 0.99 |
| 4541.1 | HGNC:7462 | NADH2_u10m | mitochondrially encoded NADH dehydrogenase 6 | 0.99 |
| 4694.1 | HGNC:7683 | NADH2_u10m | NADH:ubiquinone oxidoreductase subunit A1 | 0.99 |
| 4705.1 | HGNC:7684 | NADH2_u10m | NADH:ubiquinone oxidoreductase subunit A10 | 0.99 |
| 4695.1 | HGNC:7685 | NADH2_u10m | NADH:ubiquinone oxidoreductase subunit A2 | 0.99 |
| 4696.1 | HGNC:7686 | NADH2_u10m | NADH:ubiquinone oxidoreductase subunit A3 | 0.99 |
| 4697.1 | HGNC:7687 | NADH2_u10m | NDUFA4 mitochondrial complex associated | 0.99 |
| 4698.1 | HGNC:7688 | NADH2_u10m | NADH:ubiquinone oxidoreductase subunit A5 | 0.99 |
| 4700.1 | HGNC:7690 | NADH2_u10m | NADH:ubiquinone oxidoreductase subunit A6 | 0.99 |
| 4701.1 | HGNC:7691 | NADH2_u10m | NADH:ubiquinone oxidoreductase subunit A7 | 0.99 |
| 4702.1 | HGNC:7692 | NADH2_u10m | NADH:ubiquinone oxidoreductase subunit A8 | 0.99 |
| 4704.1 | HGNC:7693 | NADH2_u10m | NADH:ubiquinone oxidoreductase subunit A9 | 0.99 |
| 4706.1 | HGNC:7694 | NADH2_u10m | NADH:ubiquinone oxidoreductase subunit AB1 | 0.99 |
| 4707.1 | HGNC:7695 | NADH2_u10m | NADH:ubiquinone oxidoreductase subunit B1 | 0.99 |
| 4716.1 | HGNC:7696 | NADH2_u10m | NADH:ubiquinone oxidoreductase subunit B10 | 0.99 |
| 4708.1 | HGNC:7697 | NADH2_u10m | NADH:ubiquinone oxidoreductase subunit B2 | 0.99 |
| 4709.1 | HGNC:7698 | NADH2_u10m | NADH:ubiquinone oxidoreductase subunit B3 | 0.99 |
| 4710.1 | HGNC:7699 | NADH2_u10m | NADH:ubiquinone oxidoreductase subunit B4 | 0.99 |
| 4711.1 | HGNC:7700 | NADH2_u10m | NADH:ubiquinone oxidoreductase subunit B5 | 0.99 |
| 4712.1 | HGNC:7701 | NADH2_u10m | NADH:ubiquinone oxidoreductase subunit B6 | 0.99 |
| 4713.1 | HGNC:7702 | NADH2_u10m | NADH:ubiquinone oxidoreductase subunit B7 | 0.99 |
| 4714.1 | HGNC:7703 | NADH2_u10m | NADH:ubiquinone oxidoreductase subunit B8 | 0.99 |
| 4715.1 | HGNC:7704 | NADH2_u10m | NADH:ubiquinone oxidoreductase subunit B9 | 0.99 |
| 4717.1 | HGNC:7705 | NADH2_u10m | NADH:ubiquinone oxidoreductase subunit C1 | 0.99 |
| 4718.1 | HGNC:7706 | NADH2_u10m | NADH:ubiquinone oxidoreductase subunit C2 | 0.99 |
| 4719.1 | HGNC:7707 | NADH2_u10m | NADH:ubiquinone oxidoreductase core subunit S1 | 0.99 |
| 4720.1 | HGNC:7708 | NADH2_u10m | NADH:ubiquinone oxidoreductase core subunit S2 | 0.99 |
| 4722.1 | HGNC:7710 | NADH2_u10m | NADH:ubiquinone oxidoreductase core subunit S3 | 0.99 |
| 4724.1 | HGNC:7711 | NADH2_u10m | NADH:ubiquinone oxidoreductase subunit S4 | 0.99 |
| 4725.1 | HGNC:7712 | NADH2_u10m | NADH:ubiquinone oxidoreductase subunit S5 | 0.99 |
| 4726.1 | HGNC:7713 | NADH2_u10m | NADH:ubiquinone oxidoreductase subunit S6 | 0.99 |
| 374291.1 | HGNC:7714 | NADH2_u10m | NADH:ubiquinone oxidoreductase core subunit S7 | 0.99 |
| 4728.1 | HGNC:7715 | NADH2_u10m | NADH:ubiquinone oxidoreductase core subunit S8 | 0.99 |
| 4723.1 | HGNC:7716 | NADH2_u10m | NADH:ubiquinone oxidoreductase core subunit V1 | 0.99 |
| 4729.1 | HGNC:7717 | NADH2_u10m | NADH:ubiquinone oxidoreductase core subunit V2 | 0.99 |
| 4731.1 | HGNC:7719 | NADH2_u10m | NADH:ubiquinone oxidoreductase subunit V3 | 0.99 |
| 84701.1 | HGNC:16232 | CYOOm3 | cytochrome c oxidase subunit 4I2 | 0.99 |
| 1346.1 | HGNC:2287 | CYOOm3 | cytochrome c oxidase subunit 7A1 | 0.99 |
| 9167.1 | HGNC:2289 | CYOOm3 | cytochrome c oxidase subunit 7A2 like | 0.99 |
| 125965.1 | HGNC:24380 | CYOOm3 | cytochrome c oxidase subunit 6B2 | 0.99 |
| 4512.1 | HGNC:7419 | CYOOm3 | mitochondrially encoded cytochrome c oxidase I | 0.99 |
| 4513.1 | HGNC:7421 | CYOOm3 | mitochondrially encoded cytochrome c oxidase II | 0.99 |
| 4514.1 | HGNC:7422 | CYOOm3 | mitochondrially encoded cytochrome c oxidase III | 0.99 |

#### Supplementary table 13. Estimated parameters for the non-linear mixed effect model for the HEK293 growth data

| | Value | Std.Error | DF | t-value | p-value |
|---|---|---|---|---|---|
| **asym** | 37.0925114 | 0.79161899 | 222 | 46.8565205 | 4.36E-117 |
| **thresh** | 4.44277587 | 0.09155386 | 222 | 48.5263617 | 3.65E-120 |
| **scal.(Intercept)** | 0.95132841 | 0.07597008 | 222 | 12.5224089 | 1.44E-27 |
| **scal.backgroundMMUT** | 0.07985848 | 0.1002211 | 222 | 0.79682308 | 4.26E-01 |
| **scal.KOCPT2** | 0.02197599 | 0.14156792 | 222 | 0.15523286 | 8.77E-01 |
| **scal.KOHIBCH** | 0.52545137 | 0.1138489 | 222 | 4.61533979 | 6.64E-06 |
| **scal.KOSDHa** | -0.4084209 | 0.10191908 | 222 | -4.0073053 | 8.38E-05 |
| **scal.backgroundMMUT:KOCPT2** | -0.1426607 | 0.18047027 | 222 | -0.790494 | 4.30E-01 |
| **scal.backgroundMMUT:KOHIBCH** | -0.613164 | 0.15179324 | 222 | -4.0394684 | 7.38E-05 |
| **scal.backgroundMMUT:KOSDHa** | -0.1393123 | 0.13995724 | 222 | -0.9953917 | 3.21E-01 |

#### Supplementary table 14. Interaction contrasts estimated using non-linear mixed effect model of the HEK293 growth data

| background | KO | estimate | SE | df | t.ratio | p.value |
|---|---|---|---|---|---|---|
| MMUT - WT | CPT2 - None | -0.1426607 | 0.17685326 | 222 | -0.8066611 | 0.42072506 |
| MMUT - WT | HIBCH - None | -0.613164 | 0.14875099 | 222 | -4.1220837 | 5.30E-05 |
| MMUT - WT | SDHa - None | -0.1393123 | 0.1371522 | 222 | -1.0157494 | 0.31085445 |

#### Supplementary table 15. Parameters used for the FogBank algorithm

| **Foreground Segmentation Parameters** | |
|---|---|
| Min Cell Area | 300 |
| Fill Holes Smaller Than | 2000 |
| Fill Holes Larger Than | Inf |
| Keep Holes with Operator | AND |
| Fill Holes with percentile intensity less than | 0 |
| Fill Holes with percentile intensity larger than | 100 |
| Morphological Operation: none with radius 2 Greedy | -10 |
| **Object Separation** | |
| Min Seed Size | 150 |
| Apply Fogbank On | Grayscale Image |
| Fogbank Direction | Min -> Max |
| Min Object Area | 150 |

#### Supplementary table 16. *in silico* DMEM constraints applied

| Reaction name | Description | Lower bound | Upper bound |
|---|---|---|---|
| EX_glc(e) | D-Glucose exchange | -1 | 1000 |
| EX_gln_L(e) | exchange reaction for L-glutamine | -1 | 1000 |
| EX_arg_L(e) | L-Arginine exchange | -1 | 1000 |
| EX_Lcystin(e) | L-Cystine exchange | -1 | 1000 |
| EX_his_L(e) | exchange reaction for L-histidine | -1 | 1000 |
| EX_ile_L(e) | L-Isoleucine exchange | -1 | 1000 |
| EX_leu_L(e) | L-Leucine exchange | -1 | 1000 |
| EX_lys_L(e) | L-Lysine exchange | -1 | 1000 |
| EX_met_L(e) | L-Methionine exchange | -1 | 1000 |
| EX_phe_L(e) | exchange reaction for L-phenylalanine | -1 | 1000 |
| EX_thr_L(e) | L-Threonine exchange | -1 | 1000 |
| EX_trp_L(e) | L-Tryptophan exchange | -1 | 1000 |
| EX_tyr_L(e) | L-Tyrosine exchange | -1 | 1000 |
| EX_val_L(e) | L-Valine exchange | -1 | 1000 |
| EX_gly(e) | exchange reaction for Glycine | -1 | 1000 |
| EX_ser_L(e) | exchange reaction for L-serine | -1 | 1000 |
| EX_co2(e) | CO2 exchange | 0 | 1000 |
| EX_o2(e) | exchange reaction for oxugen | -10 | 0 |
| EX_ca2(e) | Calcium exchange | -1 | 1000 |
| EX_cl(e) | exchange reaction for Chloride | -1000 | 1000 |
| EX_fe3(e) | Fe3+ exchange | -1 | 1000 |
| EX_h(e) | exchange reaction for proton | -100 | 1000 |
| EX_h2o(e) | H2O exchange | -100 | 1000 |
| EX_k(e) | K+ exchange | -1 | 1000 |
| EX_na1(e) | exchange reaction for Sodium | -1000 | 1000 |
| EX_oh1(e) | exchange reaction for hydroxide ion | -1000 | 1000 |
| EX_pi(e) | Phosphate exchange | -100 | 1000 |
| EX_so4(e) | Sulfate exchange | -1000 | 1000 |
| EX_inost(e) | myo-Inositol exchange | -0.1 | 1000 |
| EX_pnto_R(e) | (R)-Pantothenate exchange | -1 | 1000 |
| EX_pydx(e) | exchange reaction for Pyridoxal | -0.1 | 1000 |
| EX_ribflv(e) | exchange reaction for Riboflavin | -0.1 | 1000 |
| EX_ncam(e) | Nicotinamide exchange | -1 | 1000 |
| EX_thm(e) | exchange reaction for Thiamin | -1 | 1000 |
| EX_gthrd(e) | Exchange for reduced glutathione | -1 | 1000 |
| EX_btn(e) | Exchange of biotin | -1 | 1000 |
| EX_hocbl(e) | Exchange of hydroxycobalamin | -1 | 1000 |
| EX_fol(e) | exchange reaction for Folate | -1 | 1000 |
| EX_chol(e) | exchange reaction for Choline | -1 | 1000 |

### Supplementary figures

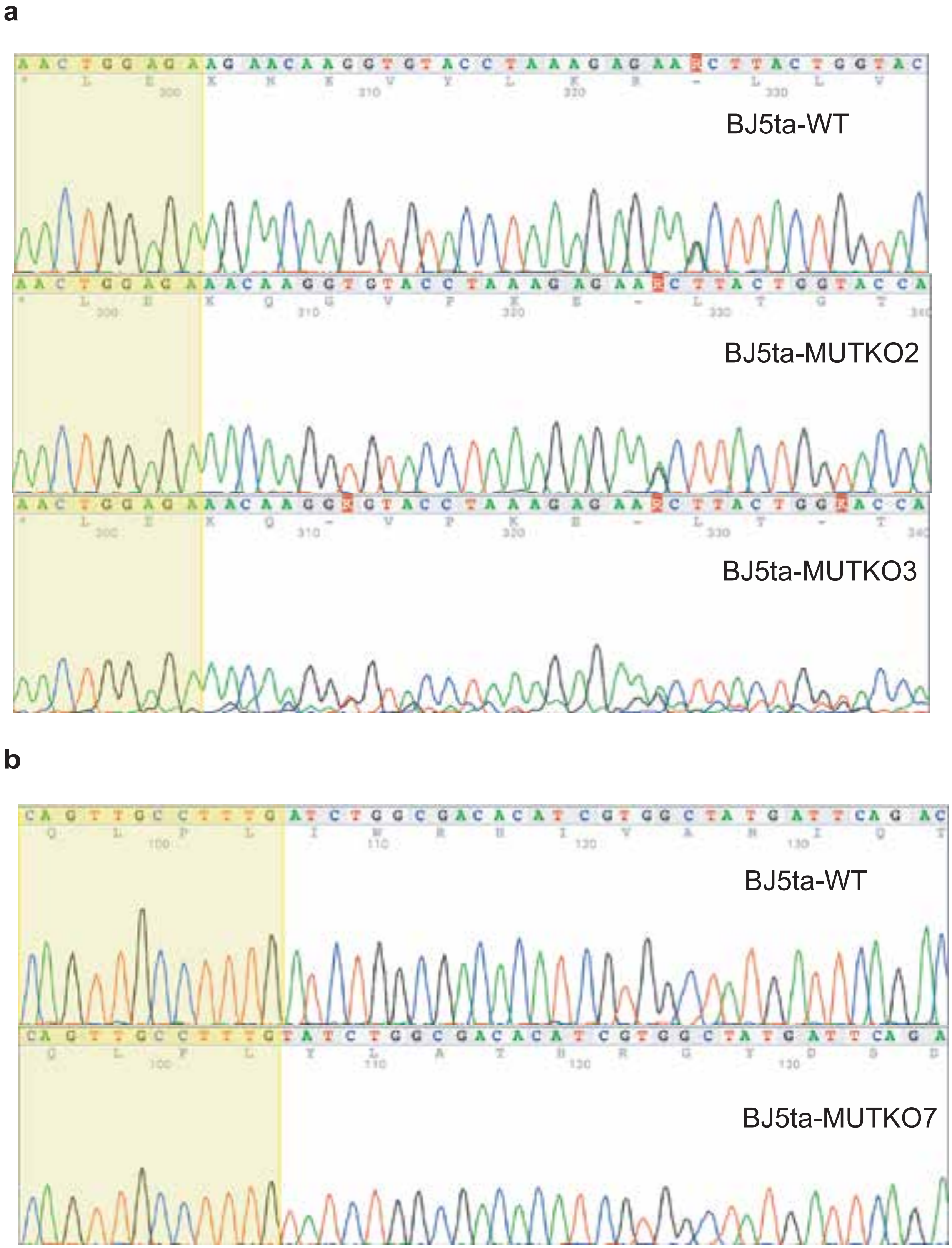


**Supplementary figure 1. Chromatographs of the mutated cell lines** - **a** Chromatographs of the cells mutated thanks to sgRNA1 vs WT cells. **b** Chromatographs of the cells mutated thanks to sgRNA2 vs WT cells.

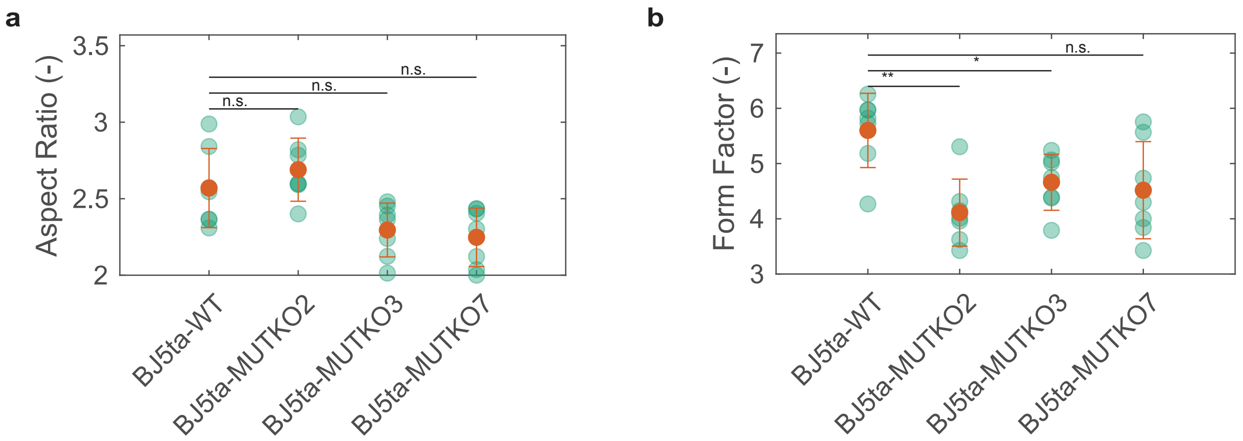


**Supplementary figure 2. Morphological characterization of mitochondria of BJ5ta fibroblasts imaged using confocal microscopy** - **a** Aspect ratio of he mitochondria for each cell line. **b** Form factor of the mitochondria for each cell line.

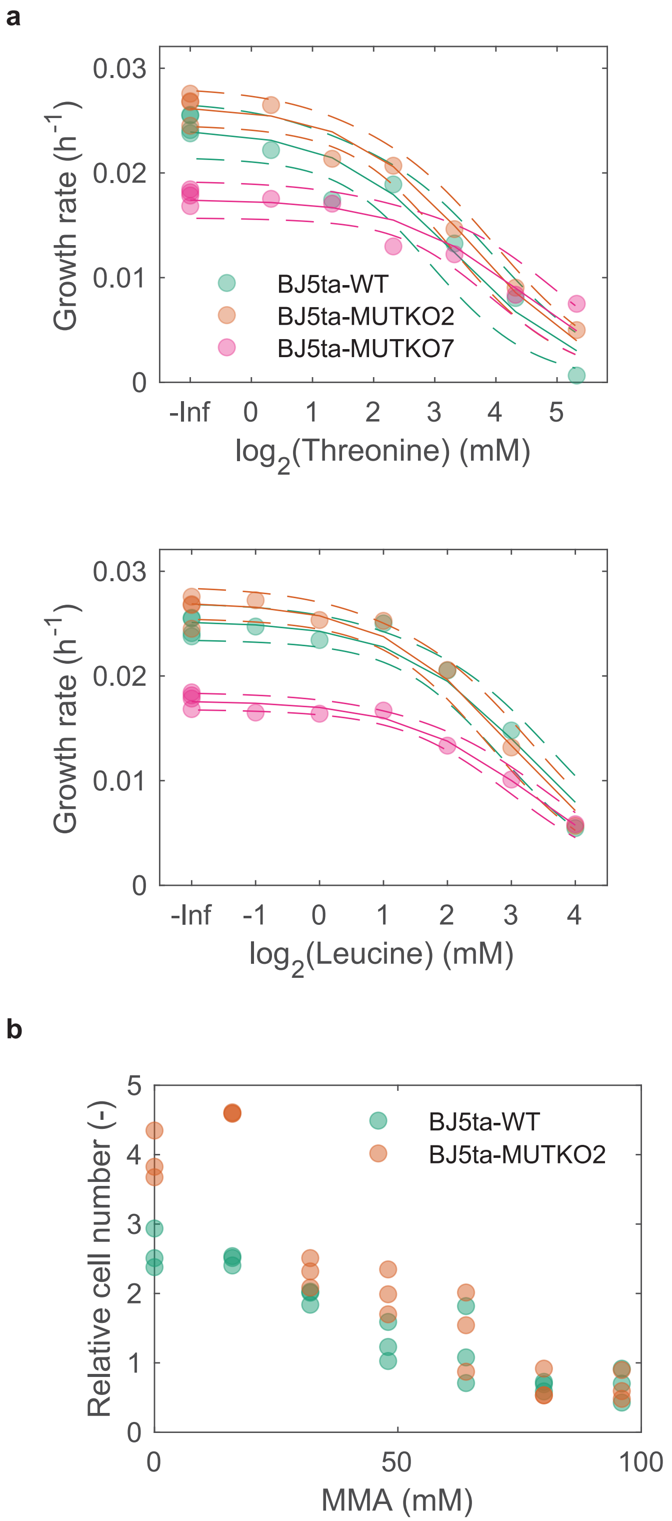


**Supplementary figure 3.** - **a** Growth rates of 3 cell lines upon addition of different doses of each amino acid Valine, Threonine, Isoleucine or Leucine. The circles are the raw data points. A 2-parameter logistic regression curve was fitted to these data points. The solid lines represent the fitted models. The dashed lines represent the 95% confidence intervals for the fitted models. **b** Total number of cells (including dead cells) divided by the initial number of cells, when cells were treated with different doses of methylmalonate (MMA).

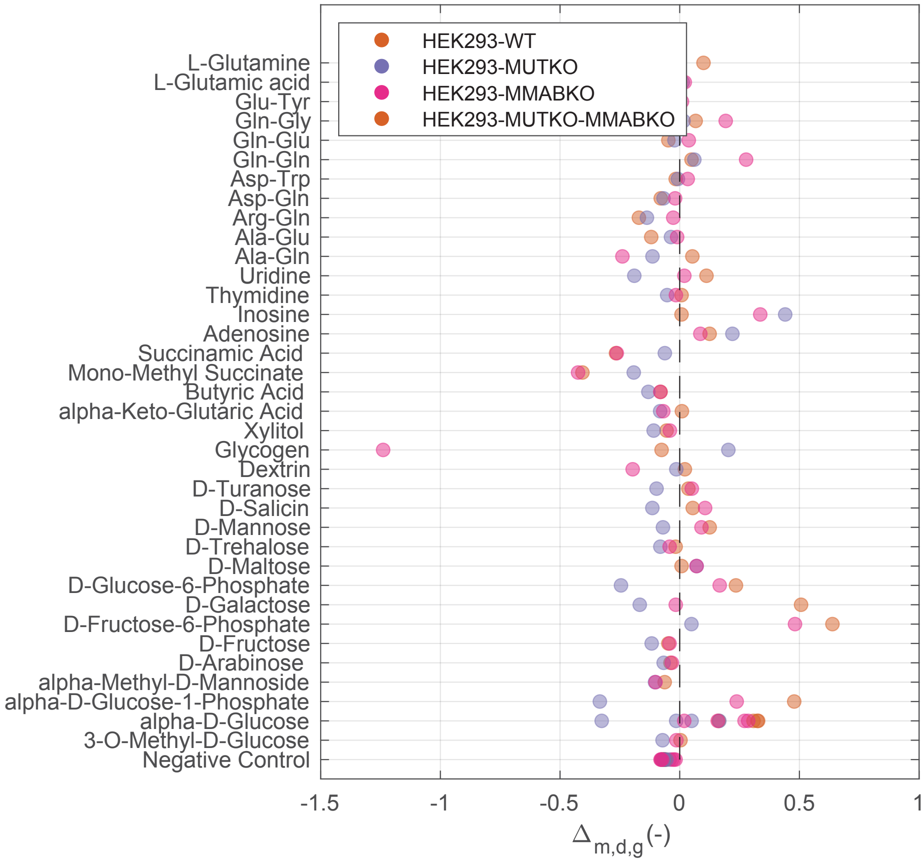


**Supplementary figure 4. Biolog phenotype microarrays** - As in Fig. 5, $\Delta_{m,d,g}$ represents the difference in absorbance for metabolite (m) in mutant (g) on day(d). Biolog phenotype microarray for different HEK293 cell lines for the metabolites selected in Lenth analysis for BJ5ta fibroblasts.

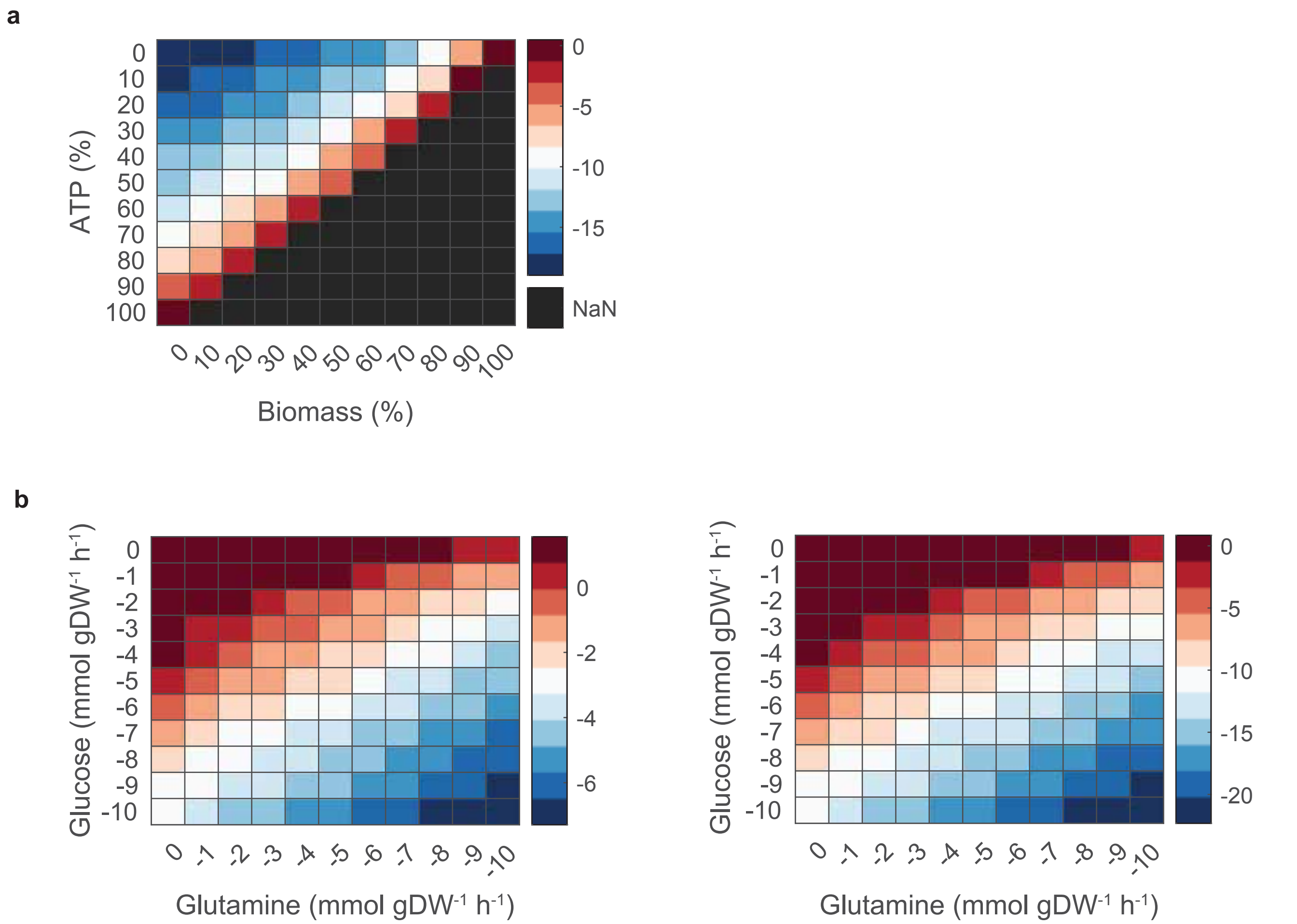


**Supplementary figure 5.** - **a** Flux variability analysis of the MMMm reaction as a function of the minimum ATP and biomass production required. Minimum flux value through the mutase reaction. **b** Flux variability analysis of the MMMm reaction as a function of the glucose and glutamine. Average of the minimum and maximum flux (left plot) and minimum flux (right plot) value through MMMm.

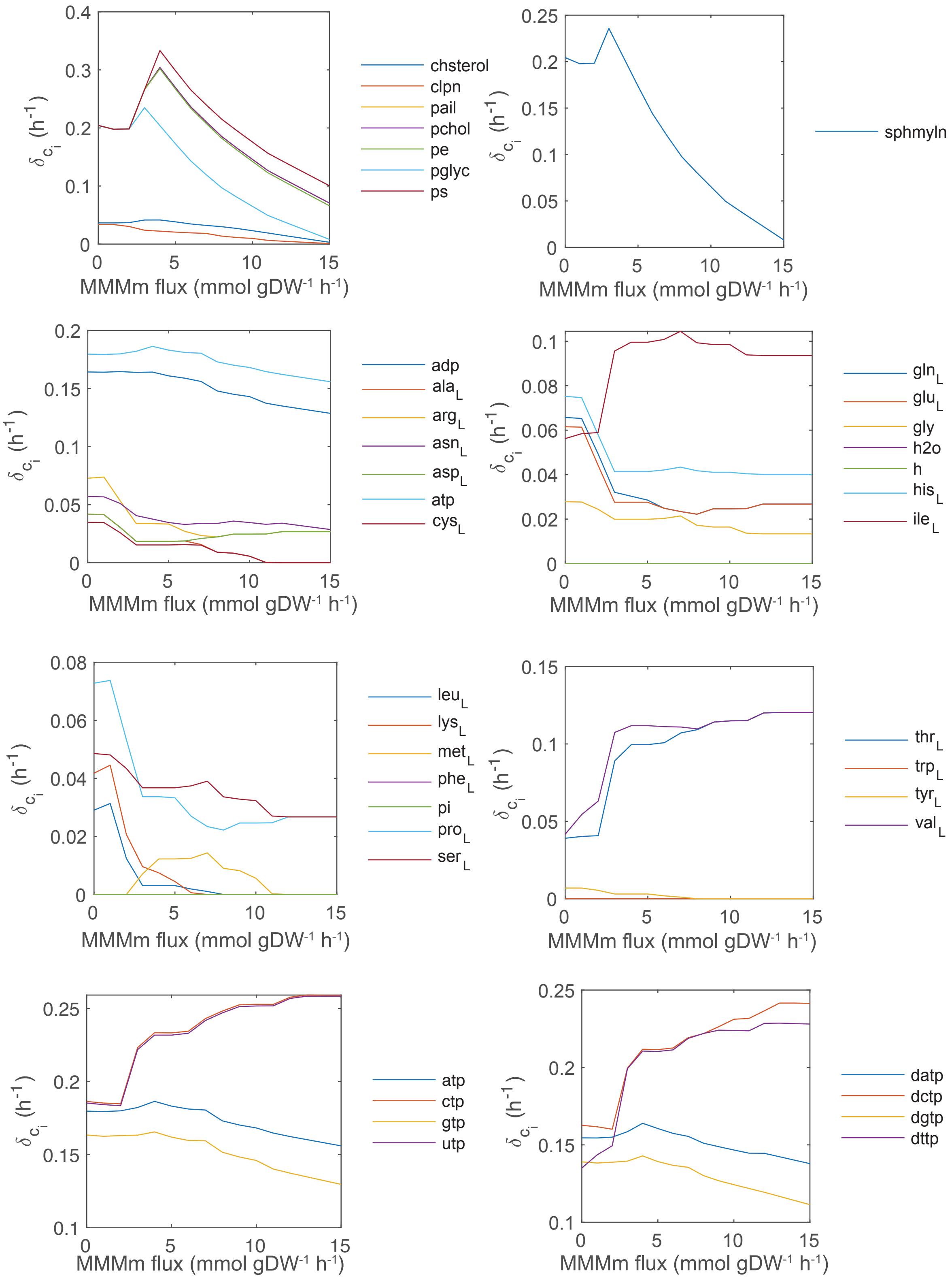


**Supplementary figure 6. Analysis of the biomass difference when adding compounds** - Biomass increase ($\delta_{c_i}$) when adding one unit flux of the compound in the legend as a function of the MMMm flux, see **Methods**.

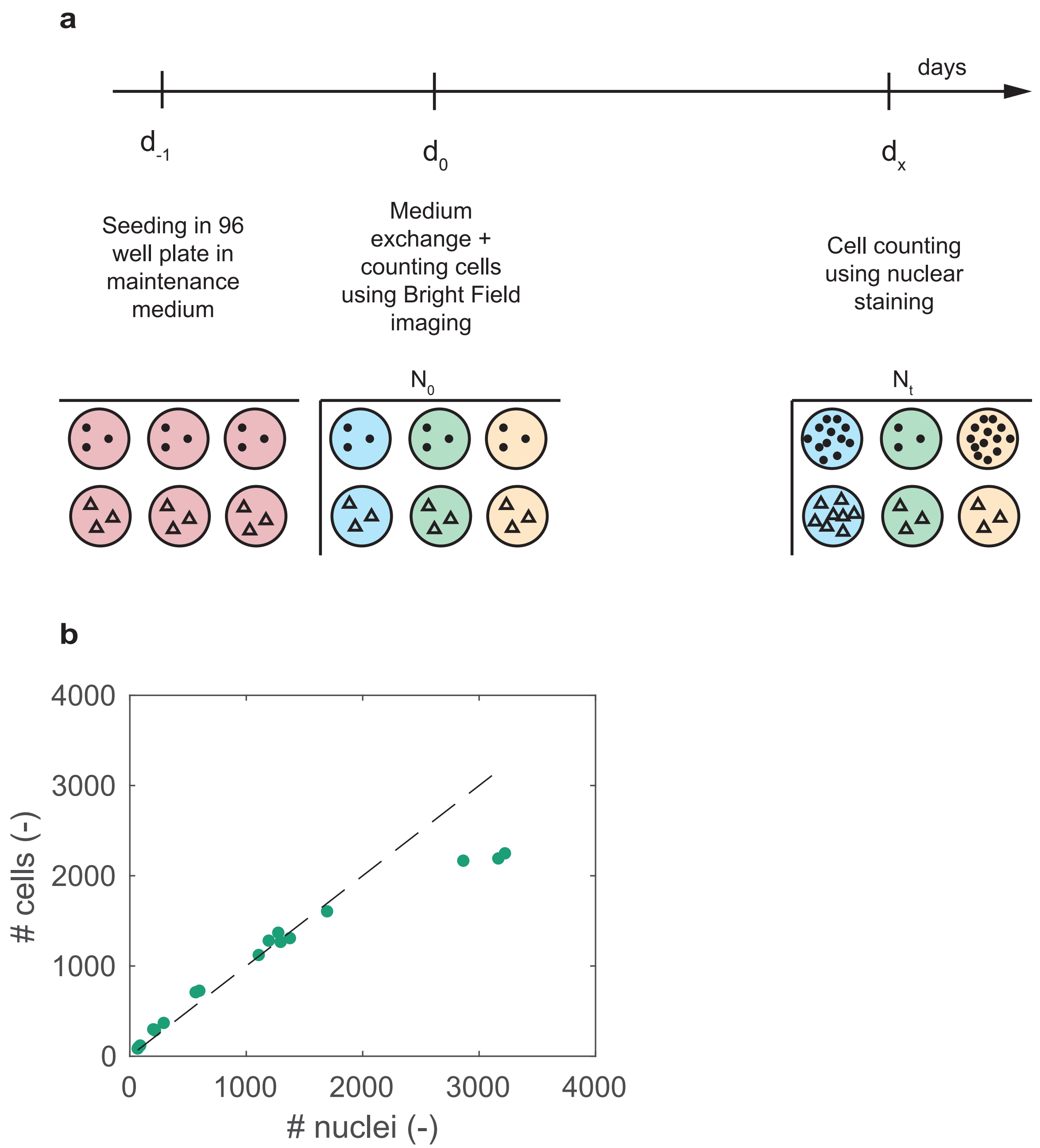


**Supplementary figure 7. Description of the growth measurement** - **a** Description of the experimental setup to measure growth rates. **b** Comparison of the number of cells in the field of view counted using nuclear staining and bright-field imaging segmentation. The dashed line is a line of slope 1.

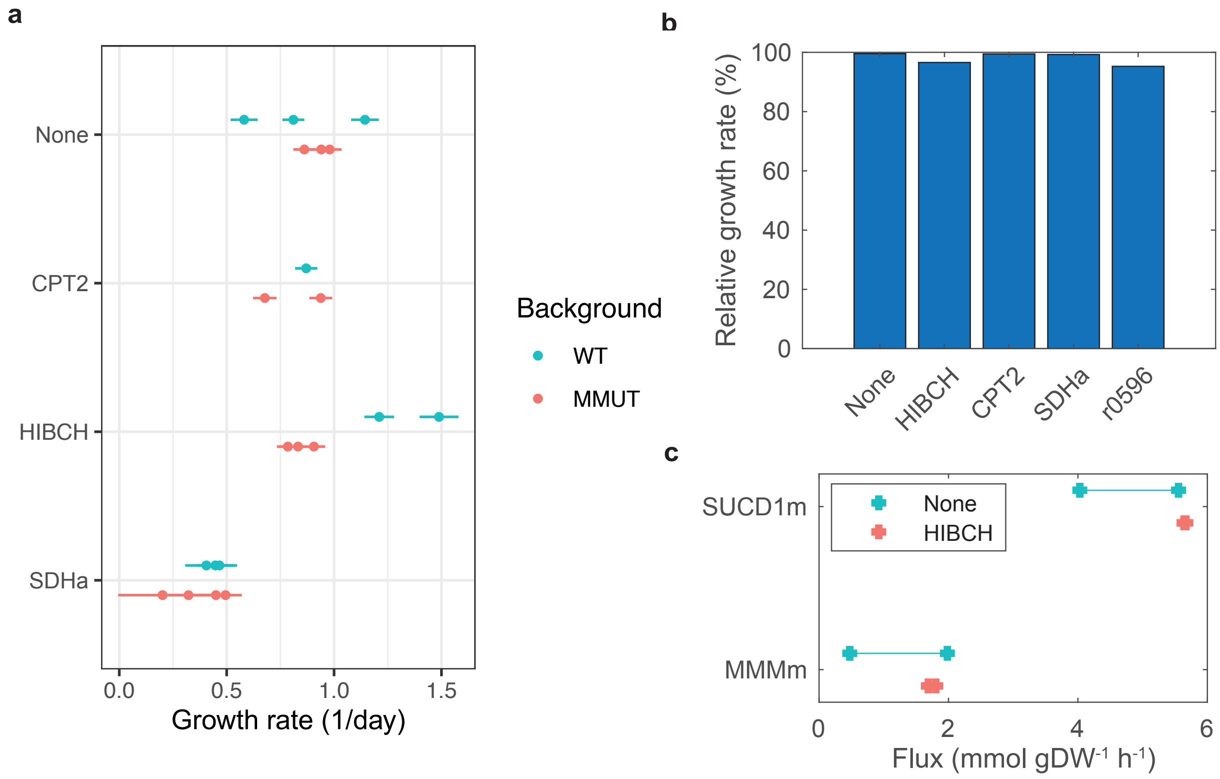


**Supplementary figure 8. Genetic interactions with *MMUT*** - **a** Growth rates of HEK293 cells with no and knockouts (*CPT2*, *HIBCH* or *SDHa*) and in either WT or *MMUT* deficient background. Growth rates were estimated using non-linear mixed effects models on the absorbance data of the Crystal Violet assay. Each dot represents the estimate for each cell line and the bars represent the uncertainty around the estimate. **b** Growth ratio of *MMUT* deficient versus WT cells as predicted using flux balance analysis. **c** Flux variability analysis for the MMMm and SUCD1m reactions for WT and *HIBCH* knock-out cells.
